# Repression of mRNA translation initiation by GIGYF1 via blocking the eIF3-eIF4G1 interaction

**DOI:** 10.1101/2023.10.14.562322

**Authors:** Jung-Hyun Choi, Jun Luo, Geoffrey G. Hesketh, Shuyue Guo, Angelos Pistofidis, Reese Jalal Ladak, Yuxin An, Tommy Alain, T. Martin Schmeing, Anne-Claude Gingras, Thomas Duchaine, Xu Zhang, Nahum Sonenberg, Seyed Mehdi Jafarnejad

**Author notes:** These authors contributed equally to this work (Jung-Hyun Choi, Jun Luo). Correspondence should be addressed to X. Z., N.S., and S.M.J.

## Abstract

Viruses commonly interfere with the function of the eukaryotic translation initiation factor 4G1 (eIF4G1), a pivotal factor in the recruitment of the eIF3 complex and ribosome to the mRNA. This results in the inhibition of general host protein synthesis and redirecting ribosomes toward viral mRNAs. Certain viruses also selectively repress the translation of mRNAs involved in the host antiviral response. GIGYF2 and its interacting cap-binding protein 4EHP enable the transcript-specific repression of mRNA translation mediated by microRNAs and RNA-binding proteins (RBPs). RNA viruses, such as SARS-CoV-2, exploit the GIGYF2/4EHP complex to selectively repress the translation of transcripts such as *Ifnb1* mRNA, which encodes the antiviral cytokine Interferon β (IFN-β). Herein, we reveal that GIGYF1, a paralogue of GIGYF2, robustly represses cellular mRNA translation through a distinct mechanism independent of 4EHP. Upon recruitment to a target mRNA by RBPs, the C-terminal region of GIGYF1 binds to subunits of eIF3 at the interaction interface of eIF3-eIF4G1. This disrupts binding of eIF3 to eIF4G1, resulting in mRNA-specific translational repression. This mechanism exerts profound influences on the host cell’s response to viral infection. Depletion of GIGYF1 induces a robust immune response by derepressing *Ifnb1* mRNA translation. Overall, our study highlights a unique mechanism of translational regulation by GIGYF1 that involves sequestering eIF3 and abrogating its binding to eIF4G1. This mechanism can be utilized by RBPs that interact with GIGYF1 to specifically repress the translation of their target mRNAs, significantly affecting critical biological processes, including host-pathogen interactions.

## Introduction

In eukaryotes, cap-dependent mRNA translation initiation involves the initial recognition of the 5’ m^7^G cap structure by a multiprotein complex known as eIF4F, which consists of the cap-binding protein eIF4E, the RNA helicase eIF4A, and the scaffold protein eIF4G1. Following cap recognition, the large eIF3 complex bridges eIF4G1 and the pre-assembled preinitiation complex (PIC), which includes the 40S ribosomal subunit, eIF1, eIF1A, and eIF2-GTP-Met-tRNAi (Hinnebusch et al., 2016). PIC scans along the 5’ UnTranslated Region (5’ UTR) of the mRNA until it reaches the translation start codon (typically AUG). This leads to the recruitment of the 60S ribosomal subunit and the subsequent assembly and precise positioning of the 80S ribosome at the start codon, thus the commencement of the polypeptide synthesis (Hinnebusch et al., 2016).

In mammals, the eIF3 complex consists of 13 subunits (eIF3A-J). The core function of eIF3 in recruiting the PIC to the mRNA exhibits a high degree of conservation from budding yeast to higher eukaryotes. Nevertheless, the *S. cerevisiae* eIF3 complex, composed of five subunits (eIF3a, eIF3b, eIF3c, eIF3g, and eIF3i), is considerably smaller than the 13-subunit mammalian eIF3 complex (Valasek et al., 2017). While eIF3 plays prominent roles in various steps of mRNA translation, knockdown studies in human cells suggested that some eIF3 subunits are dispensable for general translation initiation (Wagner et al., 2014). This suggests that certain non-core subunits of mammalian eIF3 may have evolved to fulfill specialized regulatory functions. Several studies have documented both positive and negative impacts of individual eIF3 subunits on the translation of specific mRNAs (Aguero et al., 2017; Grzmil et al., 2010; Kim et al., 2007; Shah et al., 2016). For example, the eIF3D and eIF3L subunits directly bind to the 5’ m^7^G cap and regulate the cap-dependent translation initiation of specific mRNAs (Kumar et al., 2016; Lee et al., 2016). Nonetheless, our mechanistic understanding of how eIF3 regulates the translation of specific mRNAs and its potential role in the regulation of transcript-specific translation by other factors such as RNA-binding proteins (RBPs) and microRNAs remains limited.

The Grb10-interacting GYF (glycine-tyrosine-phenylalanine) proteins 1 and 2 (GIGYF1 and GIGYF2, also known as PERQ1 and PERQ2) were originally identified through a yeast two-hybrid screen for their interaction with the adaptor protein Grb10 (Giovannone et al., 2003). The GYF domain in the mid-region of GIGYF1/2, adopts a structurally conserved fold shared among a variety of proteins across diverse eukaryotic species. The GYF domain mediates interactions with proline-rich sequences (PRS) (Freund et al., 1999), allowing GIGYF1/2 to engage with several RBPs that possess one or more proline-rich sequences, including Tristetraprolin (TTP), ZNF598, and the TNRC6(A, B, C) proteins that are among the core components of the microRNA-Induced Silencing Complex (miRISC). Thereby, through interactions with these RBPs, GIGYF1/2 can be recruited to specific mRNAs. GIGYF1/2 proteins also harbor a conserved N-terminal motif (YXYXXXXLΦ), where Φ represents a hydrophobic amino acid, which enables their specific interactions with the eIF4E homologous protein, 4EHP (also known as eIF4E2), but not to eIF4E (Morita et al., 2012; Peter et al., 2017). Similar to eIF4E, 4EHP binds to m^7^G cap, albeit with lower efficiency. Yet, 4EHP does not interact with eIF4G and thereby fails to initiate the canonical cap-dependent translation initiation (Joshi et al., 2004). Thus, through binding to 4EHP, GIGYF1/2 could repress the translation of specific mRNAs to which they are recruited via their GYF:PRS-mediated interacting RBPs (e.g. TTP). However, formation of the repressor complex with 4EHP accounts for only a fraction of translational repression of the target mRNAs by GIGYF2 (Amaya Ramirez et al., 2018; Peter et al., 2017; Xu et al., 2022), highlighting the existence of an additional, 4EHP-independent mechanism of translational repression by GIGYF2. Importantly, despite several investigations into the mechanism of GIGYF2-mediated translational regulation (Amaya Ramirez et al., 2018; Hickey et al., 2020; Kryszke et al., 2016; Morita et al., 2012; Schopp et al., 2017; Sinha et al., 2020; Zinshteyn et al., 2021), the functional mechanism and biological significance of GIGYF1 remain poorly defined.

Type I interferons (IFNs α and β) play crucial roles in the innate antiviral immune system (McNab et al., 2015), initiating a signaling cascade via the Interferon-α/β receptor (IFNAR) that triggers the JAK-STAT pathway and promotes expression of interferon-stimulated genes (ISGs) with direct antiviral effects (Wang et al., 2017). However, excessive levels of IFNs can lead to autoinflammatory and autoimmune disorders (Banchereau and Pascual, 2006; Kretschmer and Lee-Kirsch, 2017; Lee-Kirsch, 2017). Consequently, maintenance of immune homeostasis necessitates a finely tuned regulatory mechanism capable of rapid production of IFNs upon detection of viral infection while prohibiting “overshooting” of the immune response. In this context, translational regulation emerges as a rapid and flexible means of controlling IFN production in response to viral infections (Hoang et al., 2023). We recently reported the involvement of the GIGYF2/4EHP complex in repressing the translation of *Ifnb1* mRNA and the subsequent restraint of IFN-β production during RNA virus infections (Xu et al., 2022; Zhang et al., 2021). We also reported that the nonstructural protein 2 (NSP2) encoded by SARS-CoV-2 enhances the GIGYF2/4EHP-mediated translational repression of *Ifnb1* mRNA, to dampen the antiviral immune response (Xu et al., 2022). Importantly, although GIGYF1 also interacts with 4EHP in a similar manner to GIGYF2, no discernible interaction between NSP2 and GIGYF1 has been observed (Davies et al., 2020; Gordon et al., 2020; Xu et al., 2022). These findings indicate the potential existence of distinct functional and biological roles for GIGYF1 and GIGYF2 proteins, which remain unexplored.

Here, we report a previously unknown mechanism of regulation of mRNA translation by GIGYF1. Our findings show that upon recruitment to an mRNA, GIGYF1 binds to subunits of eIF3 at the interface of eIF3-eIF4G interaction, impedes the recruitment of eIF3 to the eIF4F complex, and consequently represses mRNA translation initiation. Additionally, we demonstrate that this mechanism has a profound impact on the host cell response to RNA virus infection and depletion of GIGYF1 leads to a strong immune response due to de-repression of *Ifnb1* mRNA translation.

## Results

### Potent repression of IFN-β production by GIGYF1 in a 4EHP-independent manner

GIGYF1 and GIGYF2 exhibit a high degree (55%) of similarity in their amino acid sequences, particularly in their N-terminal and central regions which respectively contain the critical 4EHP-binding (YXYXXXXLΦ) and GYF motifs (Fig. 1A and Supp. Fig. 1). The two proteins, however, substantially diverge in the sequence of their C-terminal regions, with GIGYF1 containing an exclusive poly-proline (PolyP) sequence and GIGYF2 featuring stretches of glutamic Acid-rich (E) and glutamine (Q) or proline and glutamine (PQ) rich sequences (Fig. 1A and Supp. Fig. 1). While GIGYF2 and 4EHP co-stabilize each other, as previously reported (Morita et al., 2012) and indicated by the destabilization of both proteins upon CRISPR-mediated knockout of either GIGYF2 or 4EHP, depletion of GIGYF1 had no tangible impact on 4EHP expression, and vice versa (Fig. 1B). Furthermore, analyzing the distribution of the endogenous GIGYF1 and GIGYF2 among native protein complexes by size-exclusion chromatography revealed that these two proteins predominantly elute within separate fractions (Fig. 1C). GIGYF2 was primarily eluted in fractions containing complexes of very high molecular masses >660 KDa, and displayed a similar profile to 4EHP and CNOT1, which represents the core subunit of the CCR4-NOT repressor complex, consistent with previous reports that GIGYF2 interacts with subunits of CCR4-NOT complex (Amaya Ramirez et al., 2018). In contrast, GIGYF1 predominantly emerged within lower molecular mass fractions (∼660 KDa), with a different profile to GIGYF2, CNOT1 and 4EHP (Fig. 1C). These differences in elution patterns suggest a divergence in the protein complexes in which GIGYF1 and GIGYF2 participate, their mechanisms of actions, and possibly functional significance.

**Figure 1.**
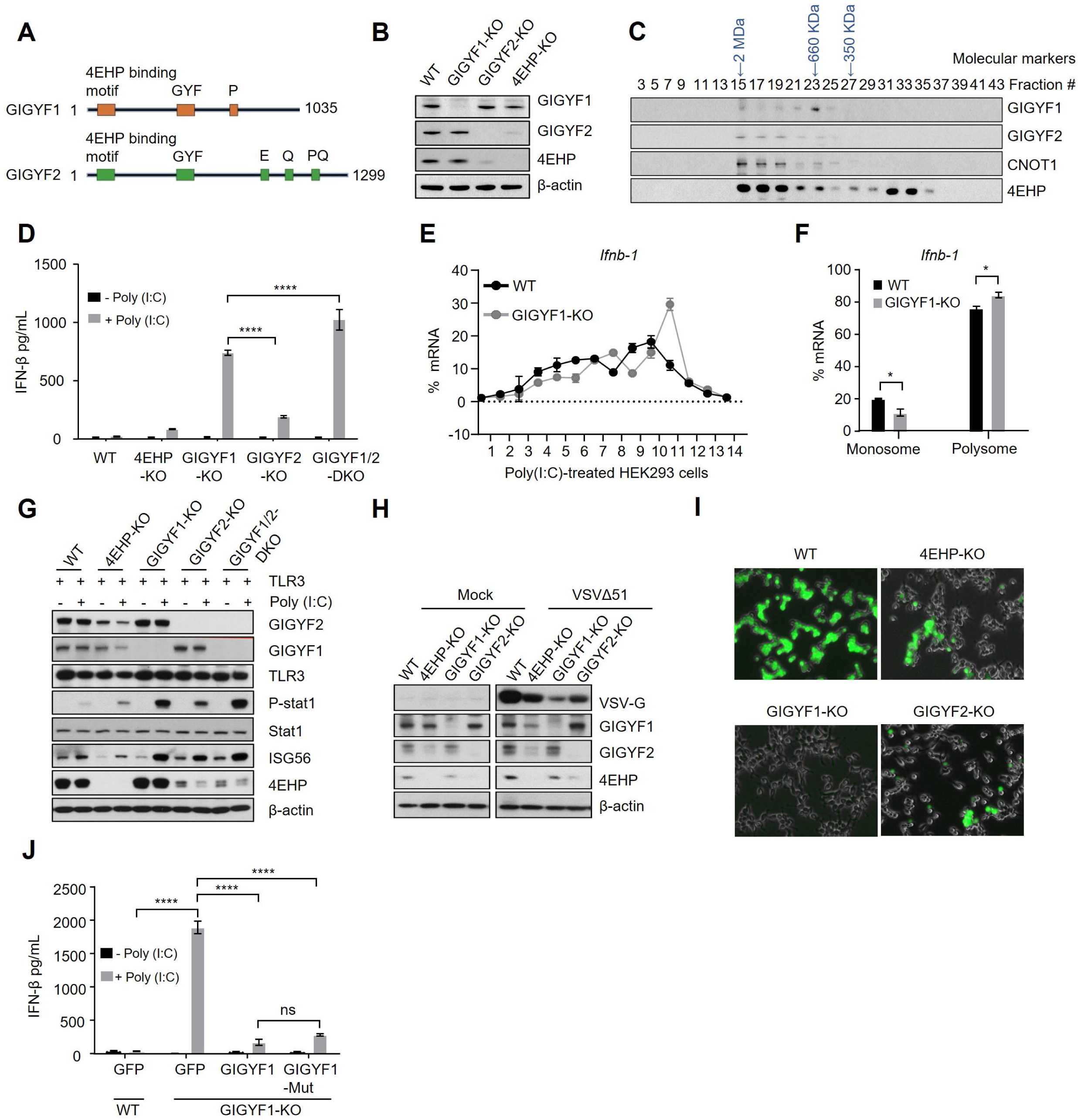
GIGYF1 enables RNA virus replication by repressing the translation of *Ifnb1* mRNA through a 4EHP-independent mechanism. (**A**) Schematic representation of the location of critical motifs in human GIGYF1 and GIGYF2 proteins. The protein length in amino acids (aa) is indicated. Domain sizes are not depicted at scale. (**B**) Western blot analysis with the indicated antibodies of WT, GIGYF1-KO, GIGYF2-KO, and 4EHP-KO HEK293 cell lysates. (**C**) Fractionation of endogenous GIGYF1, GIGYF2, CNOT1, and 4EHP by size-exclusion chromatography. A total of 10 mg of proteins from HEK293 cells was loaded onto a Superose 6 column and ran at a flow rate of 0.4 mL/min. Fractions of 0.5 mL were collected, and 50 μL of each fraction was subjected to western blot analysis. The elution position of the molecular size markers is shown. (**D**) ELISA measurement of IFN-β production in WT, 4EHP-KO, GIGYF1-KO, GIGYF2-KO, and GIGYF1/2-DKO HEK293 cells transiently expressing TLR3 following 6 h of treatment with 1 μg/mL Poly(I:C). Data are presented as mean ± SD (n=3). ****P < 0.0001; one-way ANOVA with Bonferroni’s post-hoc test. (**E**) Polysome profiling and qRT-PCR analysis of *Ifnb1* mRNA translation efficiency in poly(I:C)-treated WT and GIGYF1-KO HEK293 cells. (**F**) Relative distribution of *Ifnb1* mRNA in poorly translated polysome fractions (3 to 5) and highly translated polysome fractions (6 to 12) derived from samples described in E. Data are presented as mean ± SD. *P < 0.05; two-way ANOVA with Bonferroni’s post-hoc test. (**G**) Western blot analysis of cell lysates from (D) with the indicated antibodies. (**H** & **I**) Virus protection assay for measurement of the impact of GIGYF1 depletion on viral replication and spread. WT, 4EHP-KO, GIGYF1-KO, and GIGYF2-KO TLR3-HEK293 cells were treated with poly(I:C) for 6 h. The culture media from the treated cells were then transferred to recipient untreated TLR3-HEK293 cells, which were subsequently exposed to the VSVM51 Vesicular Stomatitis Virus at a MOI of 0.01. Virus replication was visualized by western blot with the indicated antibodies (**H**) and fluorescence microscopy (**I**). (**J**) v5-GFP control, v5-GIGYF1, or the 4EHP-binding mutant (Y39A, Y41A, M46A, L47A) v5-GIGYF1-Mut plasmids were transfected into the WT or GIGYF1-KO cells. IFN-β ELISA was performed following 6 h of 1 μg/mL HMW poly(I:C) stimulation. Data are presented as mean ± SD (n=3). ns = non-significant, ****P < 0.0001; one-way ANOVA with Bonferroni’s post-hoc test.

We previously reported that 4EHP and GIGYF2 repress the translation of the *Ifnb1* mRNA, resulting in enhanced replication of RNA viruses (Xu et al., 2022; Zhang et al., 2021). Thus, to study the molecular function of GIGYF1, we first evaluated its role in the repression of translation of *Ifnb1*, as a cellular mRNA model, and its impact on viral replication, in comparison with GIGYF2 and 4EHP. We used a HEK293 cell model that expresses the human toll-like receptor 3 (TLR3), a receptor recognizing double-stranded viral RNAs. Upon stimulating the cells with polyinosinic-polycytidylic acid [poly(I:C)], a double-stranded RNA mimic and TLR agonist, we measured IFN-β levels in wild-type (WT), 4EHP-knockout (KO), GIGYF1-KO, GIGYF2-KO, and GIGYF1/2 double knockout (GIGYF1/2-DKO) cells by ELISA. While both 4EHP-KO and GIGYF2-KO resulted in substantially increased IFN-β production (3.8 ± 0.8-fold and 8.7 ± 1.8-fold increase, respectively) compared to the WT cells, the GIGYF1-KO cells showed a comparatively more profound 33.6 ± 6.2-fold increase in IFN-β production (Fig. 1D). Notably, we observed a slightly augmented increase in IFN-β production in GIGYF1/2-DKO cells, compared with GIGYF1-KO cells. This additional increase further highlights the divergent mechanisms by which GIGYF1 and GIGYF2 repress the translation of their common target mRNAs. Measurement of mRNA expression revealed no significant alternations in *Ifnb1* mRNA levels among any of the indicated genotypes (Supp. Fig. 2A), suggesting that the observed increased IFN-β production is not due to changes in mRNA expression. A similar exacerbated increase in IFN-β expression in GIGYF1-KO cells, compared with their GIGYF2-KO, 4EHP-KO, and WT counterparts was also observed in A549 human lung epithelial cells (Supp. Fig. 2B-C). To determine the impact of GIGYF1 depletion on the translation of *Ifnb1* mRNA, we performed a polysome profiling assay using poly(I:C)-treated WT and GIGYF1-KO HEK293 cells. GIGYF1-KO did not affect the global mRNA translation, as evidenced by similar polysome/monosome ratios between the WT and GIGYF1-KO cells in both vehicle– and poly(I:C)-treated conditions (Supp. Fig. 2D-E). However, *Ifnb1* mRNA exhibited a significant shift towards heavier fractions in poly(I:C)-treated GIGYF1-KO cells compared to WT cells (Fig. 1E-F). These findings provide compelling evidence supporting the potent role of GIGYF1 in repressing the translation of *Ifnb1* mRNA.

We proceeded to explore the biological significance of the increased IFN-β production in poly(I:C)-treated GIGYF1-KO cells by measuring the activity of the STAT1 signaling pathway downstream of IFN-α/β receptor IFNAR. Western blot analysis showed enhanced phosphorylation of STAT1 and increased expression of ISG56 in 4EHP-KO and GIGYF2-KO, compared with the WT cells. Notably, GIGYF1-KO cells exhibited a further exacerbated expression of both phospho-STAT1 and ISG56 (Fig. 1G). Next, we investigated whether the GIGYF1-mediated repression of IFN-β production could effectively suppress viral replication and dissemination. To assess this, we employed a virus protection assay, a method that evaluates the ability of culture media derived from cells stimulated with poly(I:C) to protect recipient cell populations against virus infection upon transfer of the culture medium (Alain et al., 2010). We treated TLR3-HEK293 cells of the WT, 4EHP-KO, GIGYF1-KO, and GIGYF2-KO genotypes with poly(I:C) for 6 hours. Following this, we transferred the culture media from the treated cells to recipient TLR3-HEK293 cells that were left untreated. These recipient cells, referred to as “primed” cells, were subsequently exposed to a GFP-expressing Vesicular Stomatitis Virus (VSV) harboring mutations in the M protein (VSVΔ51). Viral replication levels were assessed by measuring the expression of the viral VSV-G protein with western blot and GFP with fluorescence imaging. Consistent with the elevated levels of IFN-β, phospho-STAT1, and ISG56 expression, both western blot (Fig. 1H) and fluorescence imaging (Fig. 1I) showed substantially lower levels of viral protein expression in GIGYF1-KO, compared with the 4EHP-KO and GIGYF2-KO, as well as WT cells. These data underscore that GIGYF1 potently represses IFN-β production, even more efficiently than GIGYF2 and 4EHP, and emphasize that the GIGYF1-mediated repression of IFN-β production is critical for efficient RNA virus replication.

We next examined the role of 4EHP in GIGYF1-mediated robust repression of IFN-β production by generating a mutated isoform of GIGYF1 (GIGYF1-Mut) which cannot bind to 4EHP, as confirmed by co-immunoprecipitation assay (Supp. Fig. 2F). We complemented the GIGYF1-KO HEK293 cells with either the wild-type or GIGYF1-Mut constructs (Supp. Fig. 2G). The 4EHP-binding deficient GIGYF1 very effectively, albeit to a slightly lesser extent than the wild-type isoform, repressed IFN-β production compared with the GIGYF1-KO cells that expressed GFP as a control (Fig. 1J). This indicates that, in contrast to GIGYF2-mediated repression of *Ifnb1* mRNA that largely relies on 4EHP (Xu et al., 2022), GIGYF1-mediated repression is mainly through a mechanism independent of 4EHP.

### The C-terminal region of GIGYF1 is required for translational repression of target mRNAs

Having established the ability of GIGYF1 to efficiently repress mRNA translation and its important biological role in the regulation of the host cell response to RNA virus infection, we set out to discover the mechanism underlying the efficient and 4EHP-independent translational repression by GIGYF1. 3’ UTRs serve as primary binding sites for microRNAs and RBPs, such as TTP, which recruit GIGYF1/2 to the target mRNAs (Mayya and Duchaine, 2019). We previously showed that the 3’ UTR of the *Ifnb1* mRNA mediates the 4EHP-dependent as well as 4EHP-independent translational repression by GIGYF2 (Xu et al., 2022; Zhang et al., 2021). Therefore, as an experimental model, we used a reporter system wherein a Renilla Luciferase (RL) open-reading frame was fused to the 3’ UTR derived from the human *Ifnb1* mRNA. Compared to the control reporter, the *Ifnb1* 3’ UTR construct was repressed by 37 ± 5% in WT cells (Fig. 2A). Interestingly, in the basal untreated condition, GIGYF1 knockout did not affect the repression induced by *Ifnb1* 3’ UTR (Fig. 2A). However, while poly(I:C) stimulation increased the repression of the *Ifnb1* 3’ UTR to 54 ± 9% in the WT cells, this repression was significantly relieved (24 ± 11%, p<0.01; Fig. 2A) in GIGYF1-KO cells. This contrasts with the de-repression of the *Ifnb1* 3’ UTR construct in untreated conditions (Supp. Fig. 3A), as well as poly(I:C)-treated 4EHP-KO and GIGYF2-KO cells (Xu et al., 2022). These data suggest a context-dependent mechanism governing the activation of GIGYF1-mediated repression of IFN-β production that is distinct from the GIGYF2/4EHP-mediated repression.

**Figure 2.**
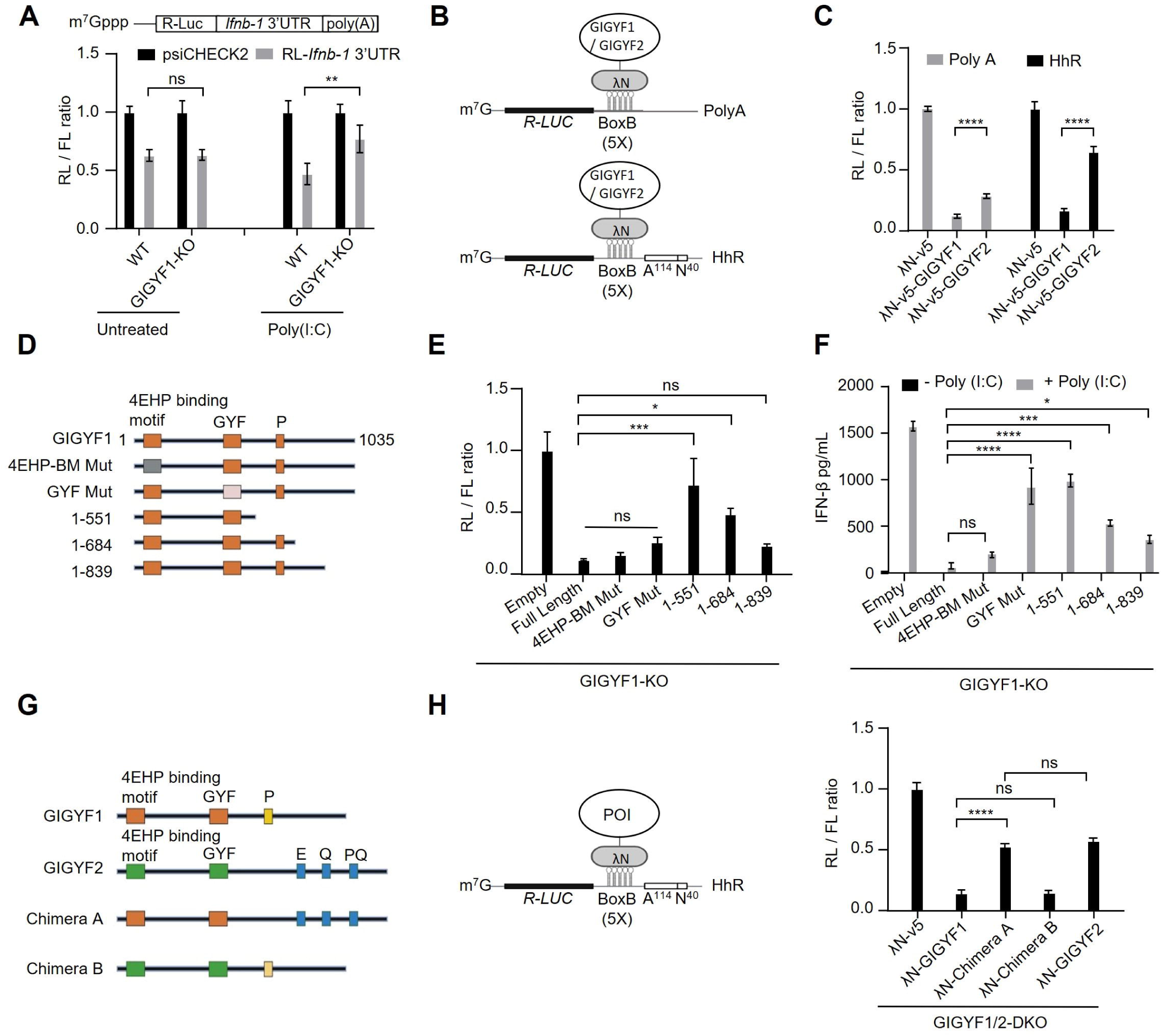
The C-terminal region of GIGYF1 is required for translational repression of target mRNAs. (**A**) *Top*: Schematic representation of the psiCHECK2-RL-*Ifnb1* 3’ UTR reporter. *Bottom*: WT and GIGYF1-KO HEK293 cells were co-transfected with psiCHECK2-RL-*Ifnb1* 3’ UTR reporter or the psiCHECK2-RL reporter without the *Ifnb1* 3’ UTR and a Firefly Luciferase (FL) reporter (as control). 18 h after transfection, cells were mock-treated or stimulated with 0.05 μg/mL HMW Poly(I:C) for 12 h, after which Renilla Luciferase (RL) and FL activities were measured. RL values were normalized against FL values, and the ratios were calculated for the psiCHECK2-RL-*Ifnb1* 3’ UTR reporter relative to the control psiCHECK2 reporter level for each condition. Data are presented as mean ± SD (n=3). ns = non-significant, **P < 0.01; two-way ANOVA with Bonferroni’s post-hoc test. (**B**) Schematic representation of the λN:BoxB tether-function system with the deadenylation-permissive RL-5boxB-polyA and deadenylation-resistant RL-5boxB-HhR (hammerhead ribozyme) reporters. (**C**) Analysis of the relative silencing of the RL-5boxB-polyA and RL-5boxB-HhR reporters upon tethering with λN-v5-GIGYF1 or λN-v5-GIGYF2 in HEK293 cells. Data are presented as mean ± SD (n=4). ****P < 0.0001; two-way ANOVA with Bonferroni’s post-hoc test. (**D**) Schematic of the domain structures of full-length GIGYF1, 4EHP-binding mutant GIGYF1, the GYF motif mutant GIGYF1, and GIGYF1 truncation mutants used in (E) and (F). Numbers refer to the position of amino acids. (**E**) Tether-function assay with the full-length GIGYF1 and the indicated mutants in GIGYF1-KO HEK293 cells. Cells were co-transfected with the indicated plasmid along with the deadenylation-resistant RL-5BoxB-HhR and FL control plasmid, followed by dual-luciferase measurement assay 24 h post-transfection. Data are presented as mean ± SD (n=3). ns = non-significant, *P < 0.05, ***P < 0.001, ****P < 0.0001; one-way ANOVA with Bonferroni’s post-hoc test. (**F**) ELISA measurement of IFN-β production in GIGYF1-KO HEK293 cells overexpressing the full-length or indicated mutant isoforms of GIGYF1, followed by 6 h treatment with 1 μg/mL HMW poly(I:C). Data are presented as mean ± SD (n=3). ns = non-significant, *P < 0.05, ***P < 0.001, ****P < 0.0001; one-way ANOVA with Bonferroni’s post-hoc test. (**G**) Schematic of the domain structures of wild type GIGYF1 and GIGYF2 and two chimeric constructs derived from the N-terminus of GIGYF1 and C-terminus of GIGYF2 wild types (Chimera A) or N-terminus of GIGYF2 and C-terminus of GIGYF1 wild types (Chimera B). (**H**) Tether-function assay for measurement of repression of the deadenylation-resistant RL-5BoxB-HhR reporter with the indicated constructs in GIGYF1/2-DKO HEK293 cells, 24 h post-transfection. Data are presented as mean ± SD (n=3). ns = non-significant, ****P < 0.0001; one-way ANOVA with Bonferroni’s post-hoc test.

To gain further mechanistic insights into the robust translational repression by GIGYF1, we employed a tether-function assay using the λN:BoxB system (Baron-Benhamou et al., 2004). Briefly, we used a Renilla Luciferase (RL) reporter containing five BoxB hairpin motifs in its 3′ UTR. The reporter was co-transfected into HEK293 cells along with a plasmid encoding a fusion of GIGYF1 or GIGYF2 linked to the λN peptide, which binds to the BoxB hairpins (Fig. 2B; *top*). Furthermore, to specifically dissect the impact of GIGYF1/2 on mRNA translation, we used a deadenylation-resistant variation of the RL-5BoxB reporter that encodes a self-cleaving Hammerhead ribozyme (HhR) at its 3′ end, generating an internalized Poly(A) stretch of 114 nucleotides to prevents deadenylation and subsequent degradation of the reporter mRNA (Fig. 2B; *bottom*) (Fukaya and Tomari, 2012). Despite comparable levels of expression of the λN-GIGYF1 and λN-GIGYF2 proteins (Supp. Fig. 3B), we observed significantly more repression, particularly of the degradation-resistant HhR reporter, by λN-GIGYF1 compared with λN-GIGYF2 (87.8% and 71.5% repression of the polyA and 83.3% and 35.4% repression of the HhR reporter for GIGYF1 and GIGYF2, respectively; Fig. 2C). We also assessed the kinetics of repression of the RL-5BoxB-polyA and RL-5BoxB-HhR reporters by increasing amounts of λN-GIGYF1 and λN-GIGYF2. With both RL-5BoxB-polyA and RL-5BoxB-HhR reporters, λN-GIGYF1 consistently demonstrated significantly stronger repression compared to λN-GIGYF2 at all tested doses of the plasmids (Supp. Fig. 3C-D). These findings further corroborate that, compared with GIGYF2, GIGYF1 exerts greater repression on mRNA translation, irrespective of its susceptibility to deadenylation or the sequence of the target mRNA. Thus, henceforth we exclusively used the HhR system to delineate the mechanisms underlying translational repression by GIGYF1.

To identify the underlying mechanism of robust translational repression of target mRNAs by GIGYF1, we constructed a series of mutants and truncated isoforms of λN-GIGYF1 (Fig. 2D & Supp. Fig. 3E). Co-expression of these constructs with the RL-5BoxB-HhR reporter in GIGYF1-KO cells revealed that while mutating the 4EHP-binding or GYF motifs had no significant impact on λN-GIGYF1-mediated repression, a progressive truncation of the C-terminal region of GIGYF1 resulted in increasing de-repression of the reporter (Fig. 2E). We also tested the impact of the same mutations and truncated isoforms on IFN-β expression in poly(I:C) treated GIGYF1-KO cells. Similar to the outcomes of the tether-function experiment (Fig. 2E), mutating the 4EHP-binding motif did not significantly affect the repression of IFN-β production by GIGYF1 (Fig. 2F & Supp. Fig. 3F). Conversely, the progressive truncation of the C-terminal region of GIGYF1 resulted in an increasing de-repression of IFN-β production, compared with the full-length protein (Fig. 2F). However, in contrast to the tether-function assay (Fig. 2E), the GYF motif mutant GIGYF1 isoform failed to significantly repress IFN-β production (Fig. 2F). This discrepancy between the tether-function assay and the repression of IFN-β production can be attributed to the fact that the GYF motif mutant GIGYF1 is incapable of interacting with the RBPs that would otherwise recruit GIGYF1 to the *Ifnb1* mRNA, whereas in the tether-function assay recruitment of GIGYF1 protein to the RL-5BoxB-HhR mRNA is achieved by the λN-tag, rendering the GYF motif dispensable for repression of the reporter mRNA. To further establish the role of the C-terminal region of GIGYF1 in translational repression of target mRNAs, we created two chimeric constructs by swapping the C-terminal domains of GIGYF1 and 2 (Fig. 2G and Supp. Fig. 3G). Whereas the repressive activity of tethered Chimera A containing the N-terminus of GIGYF1 and C-terminus of GIGYF2 was comparable to the full-length GIGYF2, the repressive activity of tethered Chimera B, consisting of the N-terminus of GIGYF2 and C-terminus of GIGYF1, was comparable to the full-length GIGYF1 (Fig. 2H).

Altogether, these data demonstrate that the 4EHP-binding motif of GIGYF1 is largely dispensable for translational repression, the GYF motif is required for recruitment to the target mRNAs through interacting with RBPs but is dispensable for translational repression once GIGYF1 is recruited, and the C-terminal region effects the majority of translational repression, presumably by facilitating interaction with other protein(s).

### GIGYF1 interacts with subunits of eIF3 complex

To further characterize the mechanism underlying GIGYF1-mediated translational repression, we sought to identify its interacting partners using the BioID proximity interactome assay (Roux et al., 2012). We created stable HEK293 cell lines expressing GIGYF1 fused in-frame with an abortive biotin ligase BirA* (R118G) at its N– or C-terminus. We identified 247 unique high-confidence targets for GIGYF1 (FDR ≤ 1%), including the known interactors of GIGYF1; TNRC6A, B, and C, 4EHP (eIF4E2), and ZNF598 (Supp. Table 1). Unexpectedly, BioID also unveiled two subunits of the eIF3 complex (eIF3L & eIF3E) among the proximity interactors of GIGYF1. Notably, a recent high-throughput analysis of protein-protein interactions in HEK293T also identified subunits of eIF3 as interactors of GIGYF1 (Huttlin et al., 2021). These findings suggest that GIGYF1 may interact with one or more subunits of the eIF3 complex.

We verified the interaction between GIGYF1 and eIF3 using co-immunoprecipitation (co-IP) assay, which showed that GIGYF1 co-precipitated with eIF3L with much higher efficiency compared with GIGYF2 (Fig. 3A). Corroborating the co-IP results, proximity ligation assay (PLA) revealed that co-transfection of v5-GIGYF1 with FLAG-eIF3L, D, or G resulted in robust PLA signals, while little or no signals were detected in cells co-transfected with v5-GIGYF2 alongside the examined eIF3 subunits (Fig. 3B-C and Supp. Fig. 4A-D). Size exclusion chromatography analysis of the endogenous proteins revealed that the subunits of the eIF3 complex elute over several fractions, likely a reflection of its involvement with a wide range of complexes engaged in various stages of mRNA translation such as eIF4F, PIC, and reinitiating ribosomes (Erzberger et al., 2014). However, GIGYF1 mainly co-eluted with a fraction of eIF3 proteins characterized by an apparent molecular weight of ∼660 KDa (Supp. Fig. 4E). Notably, the molecular weight of the free eIF3 complex is estimated at 600-800 KDa (Erzberger et al., 2014). This likely indicates that GIGYF1 is associated with a fraction of eIF3 complexes that are not actively participating in active mRNA translation.

**Figure 3.**
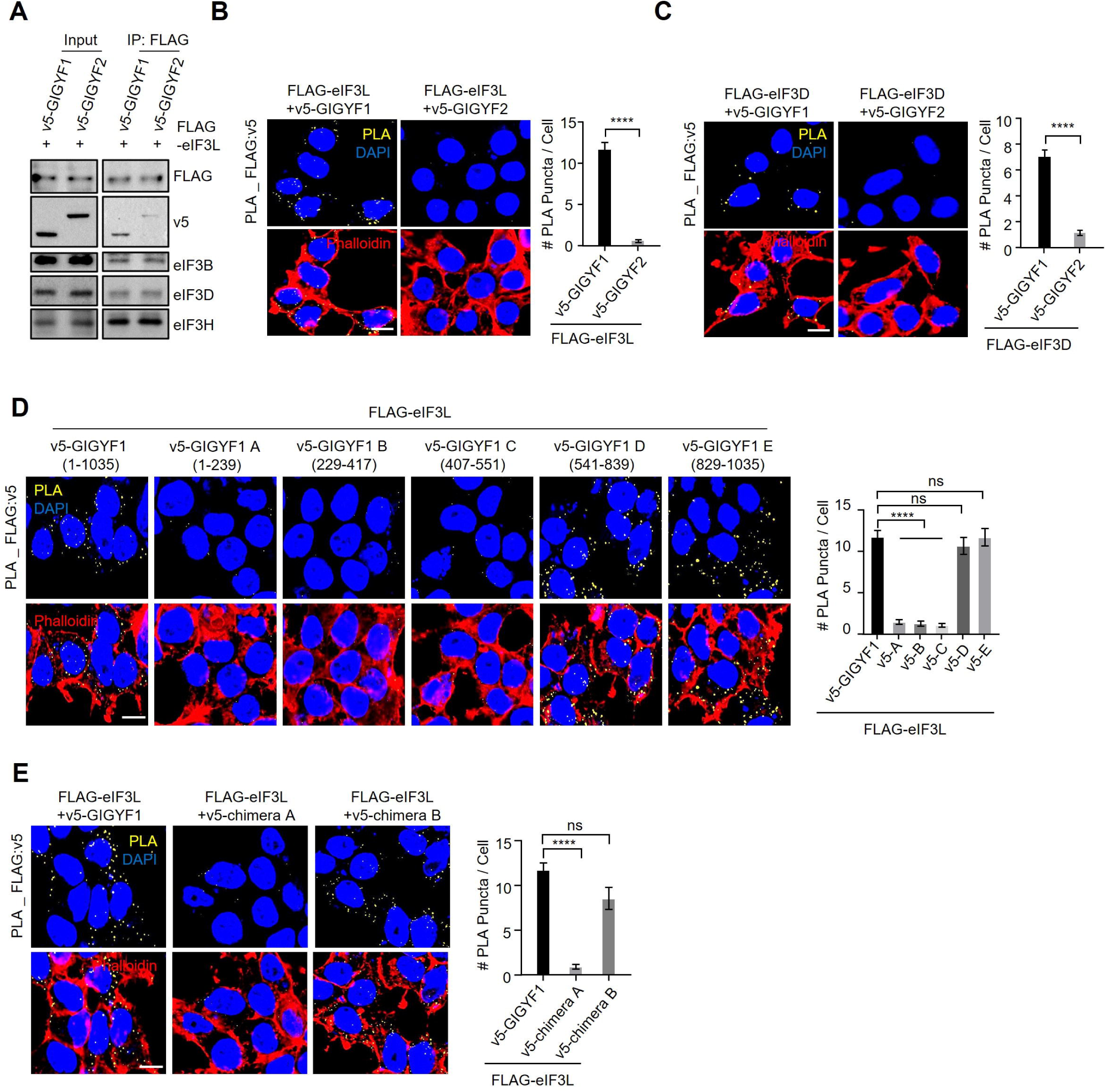
The C-terminal domain of GIGYF1 mediates its interaction with the eIF3 complex. (**A**) Co-IP assay for detection of interaction between FLAG-eIF3L and eIF3B, eIF3D, eIF3H, and v5-GIGYF1 or v5-GIGYF2 in HEK293T cells. Whole-cell lysates were prepared 24 h after transfection and subjected to immunoprecipitation using an anti-FLAG antibody followed by blotting with the indicated antibodies. (**B** & **C**) *Left*: Proximity Ligation Assay (PLA) for detection of interactions between GIGYF1 and indicated subunits of eIF3 complex. Sites of interactions are visualized as fluorescent punctate in HEK293T cells transfected with vectors expressing v5-GIGYF1 or v5-GIGYF2 together with FLAG-eIF3L (**B**) or FLAG-eIF3D (**C**). 24 h post-transfection cells were fixed and subjected to PLA using FLAG and v5 antibodies. PLA signals are shown in yellow. The nucleus and actin cytoskeleton were counterstained with DAPI and phalloidin (red), respectively. Scale bar = 10 µm. *Right*: The bar graphs represent the number of PLA signals from at least 30 cells, counted in each sample. n=5 independent experiments. ****P < 0.0001; unpaired t-test. (**D**) *Left*: HEK293T cells were co-transfected with FLAG-eIF3L and the v5-tagged full-length or indicated truncated isoforms of GIGYF1 (see Supp Fig. 4F for more details). 24 h post-transfection cells were fixed and subjected to PLA using FLAG and v5 antibodies. PLA signals are shown in yellow. The nucleus and actin cytoskeleton were counterstained with DAPI and phalloidin (red), respectively. Scale bar = 10 µm. *Right*: The bar graphs represent the number of PLA signals from at least 20 cells, counted in each sample. n=3 independent experiments. ns = non-significant, ****P < 0.0001; one-way ANOVA with Bonferroni’s post-hoc test. (**E**) *Left*: PLA assay for detecting interactions between eIF3L and the indicated chimeric constructs (described in Fig. 2G). *Right*: The bar graphs represent the number of PLA signals from at least 20 cells, counted in each sample. n=3 independent experiments. Data are presented as mean ± S.E.M. ns= non-significant, ****P < 0.0001; one-way ANOVA with Bonferroni’s post-hoc test.

To determine the specific region of GIGYF1 responsible for interaction with eIF3 subunits, we engineered five v5-tagged contiguous fragments (designated as GIGYF1 A-E; Supp. Fig. 4F) and co-transfected them with FLAG-eIF3L in HEK293T cells. PLA assay revealed interactions between eIF3L and full-length GIGYF1 as well as fragments D (541-839) and E (829-1035), both derived from the C-terminal region of GIGYF1 (Fig. 3D and Supp. Fig. 4G). Notably, while co-transfection of FLAG-eIF3L and v5-tagged Chimera A, which encompasses the N-terminal region of GIGYF1 and the C-terminal region of GIGYF2 produced only a weak signal, a strong signal was observed with Chimera B, consisting of the N-terminal of GIGYF2 and C-terminal of GIGYF1 (Fig. 3E and Supp. Fig. 4H). Similar interaction patterns were also observed for eIF3D with GIGYF1 fragments and Chimeras (Supp. Fig. 5A-D). These findings indicate that the region responsible for GIGYF1’s interaction with the eIF3 complex spans its C-terminus (residues 541-1035), which coincides with the region of the protein that is responsible for the robust repression of the target mRNAs (Fig. 2E-H).

### GIGYF1 blocks the interaction between eIF3 complex and eIF4G1

We next set out to investigate if the interactions between GIGYF1 and eIF3 play a role in the translational repression of target mRNAs by GIGYF1. Our BioID, PLA, and co-IP data consistently demonstrated that GIGYF1 interacts with eIF3 through its C-terminal region (Fig. 3), which is also responsible for orchestrating potent translational repression of mRNAs upon recruitment of GIGYF1 (Fig. 2E-H). Binding of eIF3 to eIF4G is instrumental in the recruitment of PIC to the mRNA 5’ cap. Previous structural analyses of the interactions between the eIF3 complex and eIF4G have revealed that the E, D, and C subunits of eIF3 constitute the interface through which it binds to eIF4G (Fig. 4A; (Villa et al., 2013)). Therefore, we postulated that GIGYF1 represses the translation of its target mRNAs by interacting with eIF3 at the interface where it binds to eIF4G, effectively blocking the critical eIF3-eIF4G interaction. This disruption could impede the recruitment of the PIC to eIF4F on the mRNA cap, subsequently hindering mRNA translation initiation. To test this hypothesis, we first examined the impact of GIGYF1 on interactions between subunits of eIF3 and eIF4G by co-IP. Restoring GIGYF1 expression in GIGYF1-KO HEK293 cells reduced the co-precipitation of eIF4G1, but not eIF3E, by eIF3D (Fig. 4B). Importantly, GIGYF1 overexpression did not impact upon eIF3D mediate pulldown of eIF4G2 (DAP5), a member of the eIF4G proteins family, which binds to eIF4A and eIF3, but not eIF4E, and thereby drives cap-independent translation (Henis-Korenblit et al., 2002). Furthermore, GIGYF1 overexpression did not impact upon eIF3D-mediate pulldown of eIF2α, a key subunit of PIC (Supp. Fig. 6A), suggesting the specificity of GIGYF1-mediated interruption of eIF3 interaction with eIF4G1. GIGYF1 overexpression enhanced co-precipitation of TTP with eIF3D (Supp. Fig. 6A). TTP is an RBP that is known to regulate the translation as well as stability of its target mRNAs through interactions with the GYF motifs of GIGYF1/2 (Tollenaere et al., 2019). This indicates that eIF3 could exist in a complex with RBPs that recruit GIGYF1 to repress the translation of their target mRNAs. Moreover, we observed that increasing amounts of GIGYF1 expression in GIGYF1-KO cells resulted in a reduced co-precipitation of eIF4G but not eIF3E by eIF3D (Supp. Fig. 6B). Similar results were obtained with co-IP using eIF3E, which showed a reduced association with eIF4G but not eIF3D upon expression of GIGYF1 in GIGYF1-KO cells (Supp. Fig. 6C). Overexpressing GIGYF1 also led to a diminished PLA signal between HA-eIF4G and FLAG-eIF3D, but not between HA-eIF4G and FLAG-eIF4A1, components of the eIF4F complex (Fig. 4C-D, Supp. Fig. 6D-E). These findings indicate that GIGYF1 selectively disrupts the association between the eIF3 complex and eIF4G while leaving intact the interactions between the different subunits of eIF3, PIC and eIF3, and the subunits of the eIF4F complex.

**Figure 4.**
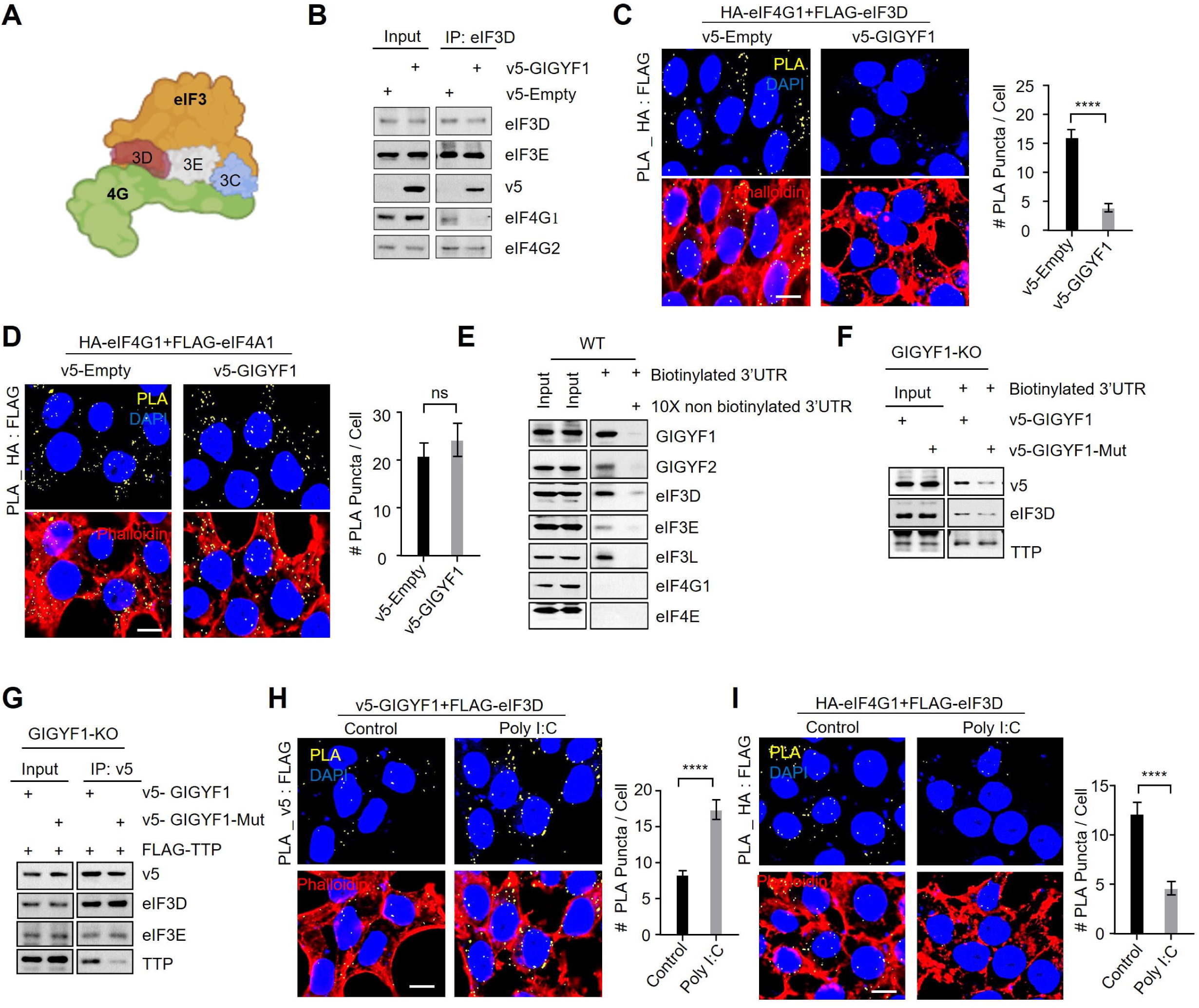
GIGYF1 blocks the interaction between eIF4G1 and eIF3 subunits. (**A**) Schematic model that highlights the subunits of eIF3 at the interface of eIF3-eIF4G1 interaction. (**B**) Co-IP assay for detection of the impact of GIGYF1 on interaction between eIF3D and eIF3E, eIF4G1, and eIF4G2. GIGYF1-KO HEK293 cells were transfected with vectors expressing v5-GIGYF1 or the v5-Empty. Whole-cell lysates were prepared 24 h after transfection and subjected to immunoprecipitation using anti-eIF3D antibody, followed by blotting with the indicated antibodies. (**C** & **D**) *Left*: PLA assay for detection of eIF4G1 and eIF3D (**C**) or eIF4G1 and eIF4A1 (**D**) interactions in GIGYF1-KO cells transfected with v5-GIGYF1 or the v5-Empty. 24 h post-transfection cells were fixed and subjected to PLA using HA and FLAG antibodies. *Right*: The bar graphs represent the number of PLA signals from at least 20 cells, counted in each sample. n=3 independent experiments. Data are presented as mean ± S.E.M. ns= non-significant, ****P < 0.0001; unpaired t-test. (**E**) Streptavidin-biotin RNA affinity purification assay with biotinylated *Ifnb1* 3’ UTR. Biotinylated *Ifnb1* 3’ UTR was incubated with lysates from HEK293 cells in the presence or absence of 10X non-biotinylated *Ifnb1* 3’ UTR for 16 h at 4°C. The pulled-down proteins were subjected to western blotting and probed with the indicated antibodies. (**F**) Biotinylated *Ifnb1* 3’ UTR was incubated with lysates from GIGYF1-KO cells expressing v5-GIGYF1 or GYF motif mutant v5-GIGYF1. The pulled-down proteins were subjected to western blotting and probed with the indicated antibodies. (**G**) Co-IP assay for detection of the impact of mutating the GYF motif on the interaction between GIGYF1, eIF3D, eIF3E, and TTP. GIGYF1-KO HEK293 cells were transfected with vectors expressing v5-GIGYF1 or GYF motif mutant v5-GIGYF1. Whole-cell lysates were prepared 24 h after transfection and subjected to immunoprecipitation using an anti-v5 antibody followed by blotting with the indicated antibodies. (**H** & **I**) *Left*: PLA assay for detection of interactions between v5-GIGYF1 and FLAG-eIF3D (H) or HA-eIF4G1 and FLAG-eIF3D (I) upon poly(I:C) stimulation. PLA signals are shown in yellow; the nucleus and actin cytoskeleton were counterstained with DAPI and phalloidin (red), respectively. Scale bar = 10 µm. *Right*: The bar graphs represent the number of PLA signals from at least 20 cells, counted in each sample. n=3 independent experiments. Data are presented as mean ± S.E.M. ns= non-significant, ****P < 0.0001; unpaired t-test.

We next queried if the interaction between GIGYF1 and eIF3 occurs within the context of the mRNAs that recruit GIGYF1 by using a streptavidin-based pulldown assay with biotinylated *Ifnb1* 3’ UTR (without cap or any associated open-reading frame). We observed that besides GIGYF1 and GIGYF2, the biotinylated *Ifnb1* 3’ UTR also precipitated eIF3L, E, and D but not eIF4G1 or eIF4E (Fig. 4E). Pulldown of these factors with biotinylated *Ifnb1* 3’ UTR was markedly reduced upon the introduction of a competitor non-biotinylated *Ifnb1* 3’ UTR (Fig. 4E), indicating the specificity of the biotinylated *Ifnb1* 3’ UTR mediated pulldown. We did not observe any visible precipitation of eIF3D or eIF3E in GIGYF1-KO cells by biotinylated *Ifnb1* 3’ UTR, despite the evident pulldown of GIGYF2 by the biotinylated *Ifnb1* 3’ UTR (Supp. Fig. 6F). This substantiates that interactions of eIF3 subunits with *Ifnb1* 3’ UTR is GIGYF1-dependent. We next examined if the GYF motif of GIGYF1, which facilitates the GYF:PRS mediated interaction with RBPs such as TTP and its recruitment to target mRNAs, is necessary for the interaction between eIF3 and target mRNAs. We observed that compared with the wild-type GIGYF1, mutation of the GYF motif led to a significant reduction in the recruitment of GIGYF1 and eIF3D to *Ifnb1* 3’ UTR (Fig. 4F). Mutating the GYF motif of GIGYF1 had no impact on the association of TTP (Fig. 4F), which is known to directly bind to the A/U-rich elements in the 3’ UTR of the *Ifnb1* mRNA (Worthington et al., 2002). Notably, mutating the GYF motif diminished the interaction between GIGYF1 and TTP, but it did not abrogate the interaction between GIGYF1 and eIF3E or D (Fig. 4G), indicating that the GYF motif is crucial for recruitment to the target mRNA, but not in the interaction with eIF3. This aligns with our previous finding that eIF3 interacts with the C-terminal of GIGYF1 (Fig. 3D-E) and that the C-terminal region, rather than the GYF domain, is responsible for the GIGYF1 mediated repression (Fig. 2E-H).

Subsequently, we investigated if the poly(I:C)-stimulated translational repression by GIGYF1 (Fig. 2A) impacts upon the interactions between eIF3 and GIGYF1 or eIF4G1. We observed a significantly augmented PLA signal between v5-GIGYF1 and FLAG-eIF3D (Fig. 4H & Supp. Fig. 6G) or between v5-GIGYF1 and FLAG-eIF3G (Supp. Fig. 6H-I) upon poly(I:C) stimulation. In contrast, poly(I:C) stimulation significantly decreased the association between eIF3D and eIF4G1 (Fig. 4I & Supp. Fig. 6J). Potential explanations for these observations are discussed below.

## Discussion

Our findings collectively support a model in which GIGYF1 inhibits mRNA translation initiation by selective disruption of the interaction between eIF3 and eIF4G1 (Fig. 5). In the last few decades, several mechanisms of regulation of general cap-dependent mRNA translation initiation have been elucidated (Tahmasebi et al., 2019). These include blocking the eIF4G and eIF4E interaction by the small eIF4E-binding proteins (4E-BPs) and inhibiting the formation of the PIC complex through phosphorylation of eIF2α. Additionally, previous studies also revealed transcript-specific mechanisms of regulation of translation initiation. For instance, repression of TOP mRNAs and microRNA target mRNAs occurs through the displacement of eIF4E from the cap by LARP1 and 4EHP, respectively (Chapat et al., 2017; Lahr et al., 2017; Naeli et al., 2023a). Furthermore, RBPs such as IRP-1 (Muckenthaler et al., 1998) and L13a (Kapasi et al., 2007) have been shown to hinder transcript-specific translation by binding to eIF4G, thereby obstructing the recruitment of the pre-initiation complex (PIC). To the best of our knowledge, this study represents the first report of a mechanism that regulates the transcript-specific translation initiation through the binding of a regulatory protein– i.e. GIGYF1-to eIF3, and specific blocking of its interaction with eIF4G1. This translational repression mechanism is transcript-specific and is likely employed by RBPs such as TTP, TNRC6s, and ZNF598 that bind to the GYF domain of GIGYF1 through their Proline-Rich Sequences (PRS). We demonstrated that this mechanism represses the translation of *Ifnb1* mRNA upon triggering the innate immune response. However, due to the broad spectrum of potential targets arising from the interactions of GIGYF1 with miRISC, ZNF598, TTP, and other RBPs, as evidenced by our BioID data (Supp. Table 1), it is expected that the specific set of cellular mRNAs regulated through the GIGYF1-eIF3 interaction mechanism is diverse and context-dependent. This is further highlighted by the observation that GIGYF1 abrogates the interaction between eIF3 and eIF4G1, but not eIF4G2 (DAP5), despite the substantial homology of DAP5 to eIF4G1 in the central segment of eIF4G1, which corresponds to the eIF4A and eIF3 binding region (Imataka et al., 1997). The distinct impacts of GIGYF1 on interactions of eIF3 with eIF4G1 and eIF4G2 is likely due to the recruitment of GIGYF1 by RBPs to mRNAs that utilize eIF4G1 mediated cap-dependent initiation, rather than the eIF4G2 mediated cap-independent initiation. Future studies employing high-throughput assays such as Ribo-Seq could unveil the comprehensive repertoire of the translatome regulated by the intricate interplay among RBPs/GIGYF1/eIF3 complexes under different conditions that induce the expression or activity of these RBPs and their target mRNAs.

**Figure 5.**
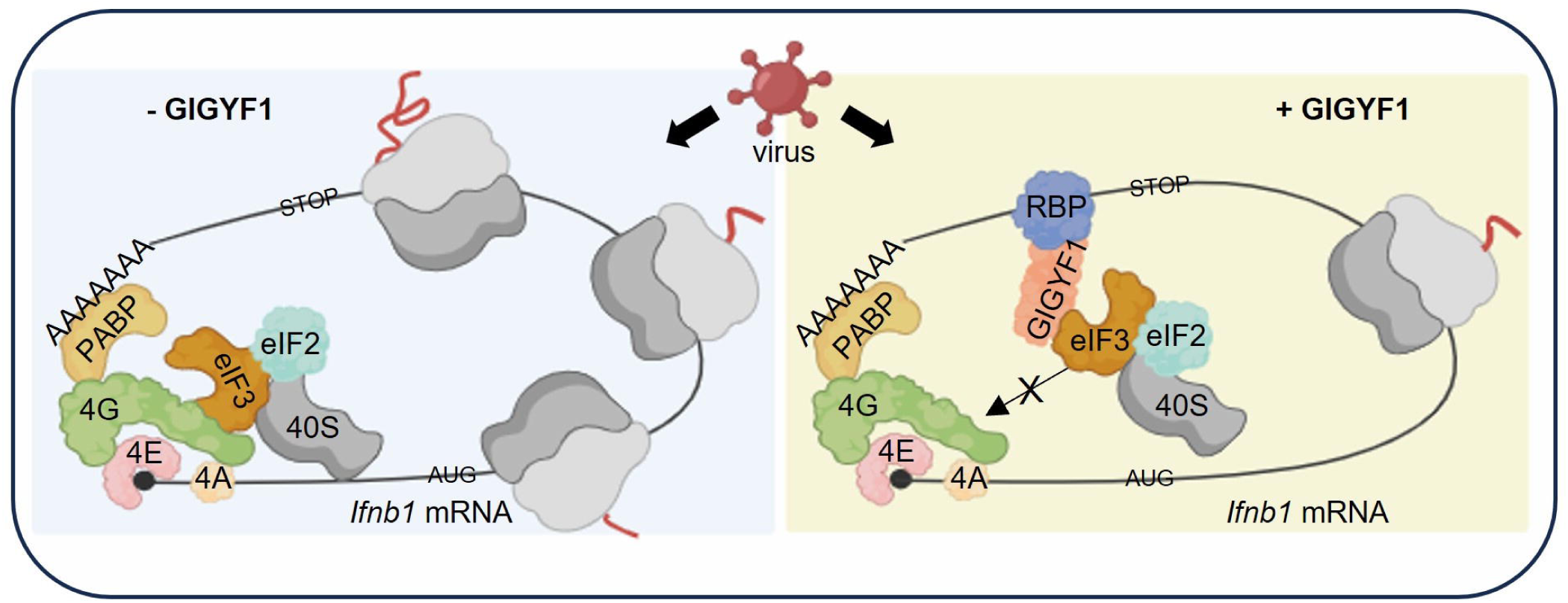
Graphic illustration of the mechanism of GIGYF1-mediated repression of the cap-dependent mRNA translation initiation. *Left*: Cap-dependent translation initiation involves the recognition of the m^7^G cap by the eIF4F complex, comprising eIF4E, eIF4A, and eIF4G1. The eIF3 complex serves as a bridge between eIF4G1 and the preinitiation complex (PIC), which includes the 40S ribosomal subunit. Upon recruitment by eIF3, the PIC scans the 5’ UTR until it reaches the translation start codon. Subsequently, it recruits the 60S ribosomal subunit, leading to the assembly of the 80S ribosome and the beginning of the elongation phase. *Right*: Upon recruitment to an mRNA by RNA-binding proteins (RBPs), typically through GYG:PRS motifs, the C-terminal region of GIGYF1 binds to the subunits of eIF3 at the interface of the eIF3:eIF4G interaction. This binding effectively blocks the recruitment of eIF3 and PIC to the eIF4F complex, thereby repressing mRNA translation initiation.

Our previous study (Chapat et al., 2017) suggested that similar to GIGYF1, 4EHP may also coexist within the same complex with eIF3E and eIF3L. This raises an intriguing possibility that GIGYF1 may employ a bifurcated mechanism to repress cap-dependent translation of target mRNAs; i) direct inhibition of the interactions between eIF3 and eIF4G1 and ii) utilization of 4EHP to displace eIF4E from the cap structure. The implications and relative contribution of each mode of translational repression exerted by GIGYF1 under different conditions require further investigation in future studies.

The complex between GIGYF2 and 4EHP serves as a key player in the translational repression triggered by microRNAs and RBPs (Amaya Ramirez et al., 2018; Hickey et al., 2020; Mayya et al., 2021; Morita et al., 2012; Tollenaere et al., 2019). While the mechanism by which GIGYF1 regulates mRNA translation has been poorly characterized, compelling evidence underscores that GIGYF2 can also exert substantial translational repression through mechanism(s) that operate independently of 4EHP (Amaya Ramirez et al., 2018; Xu et al., 2022). Our experimental data has affirmed that, similar to GIGYF2, GIGYF1 also interacts with 4EHP. However, in contrast to GIGYF2, which relies on its interaction with 4EHP for approximately 50% of the translational repression of target mRNAs (Xu et al., 2022), the interaction with 4EHP only contributes to a small fraction of GIGYF1-mediated translational repression (Fig. 1J & 2E-F). No empirically validated structures are available for either GIGYF1 or 2 proteins. Nevertheless, despite sharing a high degree (55%) of similarity of amino acid sequences, analysis of their predicted domain structures has revealed conspicuous differences in their C-terminal regions (Supp. Fig. 1). This may explain their differing abilities to interact with eIF3 subunits and utilize those interactions for the repression of target mRNAs. We noted that both the aa541-839 and aa829-1035 fragments of GIGYF1 are independently capable of interacting with the L and D subunits of eIF3 (Fig. 3D & Supp. Figure 5A). This suggests that GIGYF1 may possess multiple interaction sites with either the same or different subunits of eIF3. Consequently, further investigations are warranted to determine the structural characteristics of the C-terminal domains of GIGYF1 and GIGYF2 and elucidate the underlying mechanism behind the differential interactions of GIGYF1 and GIGYF2 with eIF3. These studies may also have implications in identifying potential sites for developing small molecule inhibitors or competitive peptides that disrupt the GIGYF1-eIF3 interaction. Such interventions could be valuable in modulating the immune system, particularly in scenarios where an overactive immune response leads to tissue damage.

We demonstrate that, through binding to eIF3, GIGYF1 represses the translation of *Ifnb1* mRNA, which encodes the crucial immunoregulatory cytokine IFN-β. This regulatory action effectively restrains the extent of activation of the host cell’s innate antiviral immune response to RNA virus infection. Congruently, the depletion of GIGYF1 conferred resistance to VSV virus replication, primarily due to the enhanced IFN-β production, indicating its significance in facilitating viral replication. Thus, both GIGYF1 and GIGYF2 play a role in the suppression of the innate immune system upon RNA virus infections, although through divergent mechanisms. It is interesting to note that this function of the GIGYF family appears to be evolutionarily conserved. The depletion of SMY2-type ILE-GYF domain-containing protein 1 (EXA1), the plant homolog of GIGYF1, has also been identified as a susceptibility gene for viral infections (Chen et al., 2022; Wu et al., 2017; Yusa et al., 2019).

Viruses employ various strategies to suppress the host’s cap-dependent mRNA translation and redirect the cellular ribosomes toward viral mRNAs (Hoang et al., 2018). Among the translation factors frequently exploited by viruses to inhibit host protein synthesis are eIF4G and eIF3. These encompass the degradation or cleavage of eIF4G1 (Belsham et al., 2000; Bovee et al., 1998), disruption of interactions between eIF4G1 and the poly(A) binding protein (PABP) (Piron et al., 1998), or sequestration of eIF4F subunits in cytoplasmic “viral factories” (Katsafanas and Moss, 2007). Several viral factors hinder host cap-dependent initiation by encoding proteins that bind to subunits of the eIF3 complex, thus interrupting its interactions with PIC (Daughenbaugh et al., 2003; Hashem et al., 2013; Sato et al., 2007; Yuan et al., 2020). An alternative mechanism has been reported for Sendai virus, where it boosts the production of the host interferon-stimulated genes ISG56 and ISGP54, which in turn inhibit translation initiation through binding to eIF3C and eIF3E, antagonizing eIF2-GTP-Met-tRNAi loading, and preventing the PIC recruitment to eIF3 (Guo et al., 2000; Hui et al., 2003; Terenzi et al., 2006). We previously reported that the SARS-CoV-2 derived NSP2 protein represses translation of *Ifnb1* (Xu et al., 2022) and other mRNAs (Naeli et al., 2023b) through co-opting the GIGYF2/4EHP complex. However, NSP2 does not interact with GIGYF1 (Davies et al., 2020; Gordon et al., 2020; Xu et al., 2022). Therefore, it remains to be understood whether viruses directly employ GIGYF1 to block the eIF4G1-eIF3 interaction and regulate host cell mRNA translation, either in a general or transcript-specific manner. Alternatively, viruses may co-opt this mechanism by enhancing the expression or activity of RBPs that recruit GIGYF1 for such regulatory purposes. Notably, enhanced expression of TTP has been observed upon infection with several types of viruses (Dudek et al., 2016; Esclatine et al., 2004; Liu et al., 2013; Schutt et al., 2023).

Disrupting the RBP/GIGYF1/eIF3 complex enhances the host antiviral response, as evidenced by increased IFN-β production, and reduced viral replication observed in GIGYF1-KO cells. Recent research has unveiled that GIGYF1 can undergo phosphorylation at multiple residues, including S638, located near the GYF motif (aa: 502-504), mediated by p38 and JNK kinases. S638 phosphorylation inhibits the GYF:PRS-mediated interactions between GIGYF1 and RBPs such as TTP and ZNF598 (Nordgaard et al., 2021). Importantly, both p38 and JNK kinases have been implicated in the host antiviral response (Faist et al., 2023; Mikkelsen et al., 2009; Wang et al., 2019). Conversely, excessive activation of p38 can lead to cytokine storm and tissue damage (Boccuni et al., 2022; Faist et al., 2023) and pharmacological inhibition of p38 has been shown to suppress the expression of pro-inflammatory cytokines, including IFN-β, in cells infected with SARS-CoV-2 (Faist et al., 2023). Thus, it is plausible that under resting conditions, the RBP/GIGYF1/eIF3 axis maintains mRNAs that encode the pro-inflammatory cytokines such as IFN-β in a translationally repressed state. Upon stimulation of the innate immune system, the interactions between GIGYF1 and eIF3 increase, thereby blocking the eIF4G1-eIF3 interactions (Fig. 4H & I). This is potentially due to the enhanced expression of mRNAs (e.g., *Ifnb1*) that harbor binding sites for RBPs (e.g. TTP) that recruit GIGYF1 and thereby elevate the frequency of GIGYF1-eIF3 interactions. An alternative possibility includes post-translational modifications or interactions with additional factors that further stimulate GIGYF1-eIF3 binding. Conversely, S638 phosphorylation of GIGYF1 and the subsequent disruption of its interaction with RBPs (e.g. TTP) provide a mechanism for rapid derepression of translation of mRNAs encoding antiviral proteins, such as IFN-β and mounting the antiviral immune response by the host cells. However, the potential role of the RBP/GIGYF1/eIF3 mechanism of translational control and the p38 and JNK-mediated S638 phosphorylation of GIGYF1 in maintaining the delicate balance between promoting the antiviral response and preventing an unrestrained proinflammatory reaction during viral infections necessitates further investigation in future studies.

In summary, our study reveals that GIGYF1 plays a crucial role in repressing cellular mRNA translation through a distinctive, transcript-specific mechanism. This mechanism entails GIGYF1’s recruitment to the mRNA through interactions with RBPs, followed by its binding to eIF3 subunits and displacing eIF4G1 from the eIF3 complex. Our findings also highlight the biological significance of this mechanism, as it effectively dampens the antiviral immune response and facilitates RNA virus infections. These results indicate the potential utility of this mechanism in devising effective methods to manipulate important biological processes such as antiviral immune response and autoimmune disorders.

## Materials and Methods

### Cell line, cell culture and transient transfection

HEK293T (Thermo Fisher Scientific) cells were maintained in Dulbecco’s modified Eagle medium (DMEM) (Wisent Inc., 319-005-CL) supplemented with 10% fetal bovine serum (FBS) (R&D Systems, S12450) and 1% Penicillin/Streptomycin (P/S) (Wisent Technologies). A549 (American Type Culture Collection [ATCC]), were maintained in RPMI (Wisent Inc., 350-000-CL), also supplemented with 10% FBS and 1% P/S. WT, 4EHP-knockout (KO), GIGYF1-KO, GIGYF2-KO, and GIGYF1/2-double knockout (DKO) HEK293 cells were maintained in DMEM supplemented with 10% FBS, 1% P/S, 100 μg/mL Zeocin (Thermo Fisher Scientific, R25001) and 15 μg/mL Blasticidin (Thermo Fisher Scientific, R210-01). All cells were cultured in a humidified atmosphere of 5% CO2 at 37°C.

### Antibodies and plasmids

The following antibodies were used: rabbit anti-GIGYF1(Bethyl, A304-132A), rabbit anti-GIGYF2 (Bethyl, A303-732A-M), rabbit anti-eIF4E2 (Genetex, GTX82524), mouse anti-β-actin (Sigma, A5441), rabbit anti-TLR3 (Cell Signaling, 6961), rabbit anti-phospho-STAT1 (Tyr701; Cell Signaling, 7649), rabbit anti-STAT1 (Cell Signaling, 14994), anti-ISG56, rabbit anti-VSV-G (Abcam, ab3556), mouse anti-v5 (Abcam, ab27671), mouse anti-HA (Biolegend, 901515), rabbit anti-HA (Sigma, H6908), mouse anti-FLAG (Abcam, ab49763), rabbit anti-FLAG (Sigma, F7425), rabbit anti-eIF3D (Abcam, ab155419, ab264228), rabbit anti-eIF4G1 (Cell signaling, 2858), rabbit anti-eIF3E (Abcam, ab36766), rabbit anti-eIF3L (Bethyl Laboratories, A304-754A), mouse anti-eIF4E (BD Biosciences, 610270), rabbit anti-TTP (Cell signaling, 71632), rabbit anti-eIF4G2 (Cell signaling, 3468), rabbit anti-eIF3H (Cell signaling, 3413), rabbit anti-eIF3B (Bethyl, A301-761A), rabbit anti-CNOT1 (Proteintech, 14276-1-AP). The psiCHECK-2 vector (Promega, C8021) and the psiCHECK-RL-*Ifnb1* 3’ UTR vector were described before (Zhang et al., 2021). Plasmids encoding λN-v5 tagged-GIGYF1, –GIGYF2, GIGYF1 truncations, Chimera A, and Chimera B were generated by cloning into pCI-neo-λN-v5 vector using XhoI and NotI restriction sites. The 4EHP-binding mutant GIGYF1 (GIGYF1 Mut; Y39A, Y41A, M46A, L47A), GYF motif mutant GIGYF1 (GIGYF1 GYF Mut; G502A, Y503A, F504A) plasmids were created using the QuikChange II Site-Directed mutagenesis kit (Agilent Technologies, 200523). The chimeric constructs were synthesized by GenScript (Piscataway, NJ, USA), and sub-cloned into the pCI-neo-λN-v5 vector using XhoI and NotI restriction sites. The sequence of the primers used are listed in Supp. Table 2.

### Generation of knockout cell lines by CRISPR-Cas9

The 4EHP-KO and GIGYF2-KO A549 cells, 4EHP-KO, GIGYF2-KO, and GIGYF1/2-double knockout (DKO) HEK293 cells were described previously (Xu et al., 2022; Zhang et al., 2021). The GIGYF1-KO A549 cells were generated using Lenti-Cas9-Blast (Addgene, plasmid 52962) and pLenti-CRISPRv2 (Addgene, plasmid 52961). Sequences of the small guide RNAs (sgRNAs) are listed in Supp. Table 2.

### Virus production

Lentivirus pseudovirions were produced by transfecting HEK293T cells using Lipofectamine2000 and 10 μg of Lentiviral plasmid, 6 μg psPAX2 (Addgene, plasmid 12260) and 4 μg pMD2.G (Addgene, plasmid 12259). 48 h post-transfection, supernatants were collected from which pseudovirions were purified by ultracentrifugation (32,000 rpm) for 2 h. The pelleted virus was resuspended into the DMEM medium. Virus titer was adjusted to 5 multiplicities of infection (MOI) prior to storage at –80°C.

Single-cycle replication-competent pseudovirions were generated by co-transfecting pVSV-eGFP-DG (addgene, 31842) with VSV glycoprotein (VSV-G) expressing plasmids into HEK293T cells. 48 h post-transfection, pseudovirus was purified from the cell culture supernatant by ultracentrifugation (35,000 rpm for 2 h at 4°C). The pelleted virions were suspended for future experiments.

### Virus protection assay

VSV protection assay was performed as previously described (Alain et al., 2010). WT, 4EHP-KO, GIGYF1-KO, and GIGYF2-KO TLR3-HEK293 cells were either mock-treated or transfected with 1 μg/mL Poly(I:C) (Sigma, P1530) using Lipofectamine2000. Supernatant was collected after 6 h and added to the recipient TLR3-HEK293 cells. After overnight incubation, primed cells were infected with VSVΔ51-GFP at 0.01 MOI for 12 h. Virus replication was visualized using western blot and fluorescence microscopy.

### Poly(I:C) treatment and ELISA

HEK293 and A549 cells were seeded at 50-60% confluency. Cells were treated for 6 h with the indicated concentration of either High Molecular Weight (HMW) Poly(I:C) (InvivoGen, tlrl-pic) or Poly(I:C) (Sigma, P1530) using Lipofectamine2000. IFN-β amounts in the culture supernatant were measured by the human IFN-β ELISA kit (R&D Systems, DIFNB0) according to the manufacturer’s protocol.

### Dual-Luciferase reporter assays

WT, 4EHP-KO, GIGYF1-KO, and GIGYF2-KO HEK293 cells, seeded at a density of 150,000 cells per well, were transfected with either 20 ng psiCHECK reporter (as control) or a construct of luciferase with full-length *Ifnb1* 3’ UTR (psiCHECK-RL-*Ifnb1* 3’ UTR reporter) using Lipofectamine2000. 24 h post-transfection, cells were lysed, followed by dual-luciferase assay (Promega, E1960). Renilla luciferase (RL) values were normalized against Firefly luciferase (FL) levels for each sample.

For the tether-function assays, we used the λN:BoxB tethering approach (Baron-Benhamou et al., 2004). Briefly, a Renilla Luciferase (RL) reporter containing five BoxB hairpins in its 3′ UTR, followed by a Poly(A) tail, and a variant form of the reporter, which instead of Poly(A) tail, contains a self-cleaving Hammerhead Ribozyme (HhR) at the 3’ end (Fukaya and Tomari, 2012), were used. The reporter was co-transfected with the plasmid encoding a λN peptide-fused protein of interest and a plasmid expressing Firefly Luciferase (FL). 24 h after transfection, luciferase activities were measured using dual-luciferase assay (Promega, E1960). RL values were normalized against FL levels to determine the effect of the tethered protein on the expression of the target mRNA.

### Polysome profiling

Polysome profiling was performed as previously described (Kim et al., 2021) with minor modifications. Briefly, HEK293 cells at 80%–90% confluency were treated with 100 μg/ml of Cycloheximide (CHX) for 5 min at 37°C. Subsequently, cells were washed twice with ice-cold PBS containing 100 μg/ml CHX, suspended in a hypotonic buffer (5 mM Tris-HCl pH 7.4, 1.5 mM KCl, 2.5 mM MgCl2, 100 μg/ml CHX, 200 μg/ml RNase inhibitor, 2 mM DTT), supplemented with EDTA-free protease inhibitor tablet (Roche, 04693124001) and lysed using a detergent mixture (0.5% Triton X-100 and 0.5% sodium deoxycholate). The amount of RNA in the supernatant was quantified using NanoDrop 2000 spectrophotometer (Thermo Fisher Scientific). Equal amounts of RNA were loaded onto a 10%–50% sucrose gradient containing 20 mM HEPES-KOH pH 7.6, 100 mM KCl, 5 mM MgCl2, 100 μg/ml of CHX, 10 μ/ml RNase inhibitor, supplemented with EDTA-free protease inhibitor tablet. Gradients were centrifuged at 36,000 rpm for 2 h at 4°C. The absorbance at 254 nm was measured to analyze the sedimentation of sucrose gradients using TracerDAQ software.

### RNA extraction and RT-PCR

Total RNA from each sample was extracted using easy-BLUE™ Total RNA Extraction Kit (iNtRON Biotechnology, 17061), following the manufacturer’s instructions. Reverse transcription was performed using oligo-dT (18) and SuperScript III (Invitrogen, 18080044), utilizing 1 μg of total RNA. The mRNA levels were analyzed using the CFX Connect Real-Time PCR Detection System (Bio-Rad) with the SensiFAST SYBR No-ROX Kit (Bioline) and gene-specific primers (Supp. Table 2). Quantification was carried out using the comparative Ct method.

### Immunoprecipitation

Cells were rinsed with cold PBS and harvested by scraping into lysis buffer (40 mM HEPES pH 7.5, 120 mM NaCl, 1 mM EDTA, 50 mM NaF, 0.3% CHAPS), supplemented with a complete EDTA-free protease inhibitor tablet (Roche, 04693124001) and phosphatase inhibitor cocktail (Sigma, P5726, P0044). Pre-cleared lysates containing 500 µg of protein were incubated with 1 µg of anti-FLAG, anti-HA, or anti-v5 antibody and 40 µl of Protein G agarose beads slurry (Millipore, 16-266) in the presence of RNase A (Thermo Scientific, EN0531) per immunoprecipitation (IP) overnight at 4°C. The beads were then washed three times for 10 min each with wash buffer (50 mM HEPES pH 7.5, 150 mM NaCl, 1 mM EDTA, 50 mM NaF, 0.3% CHAPS), supplemented with a complete EDTA-free protease inhibitor tablet and phosphatase inhibitor cocktail. Proteins were eluted from the beads in the SDS sample buffer.

### Proximity Ligation Assay (PLA)

Proximity ligation assay (PLA) was conducted using Duolink reagents (Sigma, DUO92101) in accordance with the manufacturer’s instructions. In brief, cells were fixed with a solution of 4% PFA-sucrose for 15 min and permeabilized using PBS containing 0.1% Triton X-100 for 15 min. Following that, the cells were blocked with Duolink blocking solution at 37°C for 1 h and incubated with primary antibodies overnight at 4°C. Prior to incubating with the PLA probe, the cells were washed with Wash Buffer A and then incubated with the probe for 1 h at 37°C, followed by a 30 min ligation step at 37°C. To amplify the PLA signals, an amplification buffer was applied for 100 min at 37°C. Cells were washed with Wash Buffer B and mounted onto slide glass for Airyscan microscopic imaging (Zeiss).

### Streptavidin-biotin RNA affinity purification assay

*Ifnb1* mRNA 3’ UTR (NM_002176.4) was synthesized using the HiScribe T7 High Yield RNA synthesis kit (NEB, E2040S) through *in vitro* transcription, incorporating biotinylated UTP (Jena Bioscience, NU-821-BIO16). Biotinylated 3’ UTR was then incubated with cytoplasmic lysates in dialysis buffer (10 mM HEPES pH 7.4, 90 mM KOAc, 1.5 mM MgOAc, 2.5 mM DTT) for 30 min at room temperature. In competition experiments, a 10-fold excess of non-biotinylated *Ifnb1* 3’ UTR was also included in the dialysis buffer with the biotinylated *Ifnb1* 3’ UTR. After 30 min, the binding mixture was subjected to streptavidin resin (Thermo Fisher Scientific, 20349) adsorption for 16 h at 4°C on a rotary mixer. Following three washes with a wash buffer (dialysis buffer containing 1% NP-40), proteins were eluted from the resin in the SDS sample buffer and analyzed by SDS-PAGE.

### Size exclusion chromatography

HEK293 cells were collected by centrifugation and resuspended in 500 μL of lysis buffer containing 50 mM Tris-HCl (pH 7.4), 150 mM NaCl, 1 mM EDTA (pH 8.0), 1% Triton X-100, supplemented with a complete EDTA-free protease inhibitor tablet (Roche, 04693124001) and phosphatase inhibitor cocktail (Sigma, P5726, P0044). The lysate was clarified by centrifugation at 20,000 × g for 5 min at 4°C and filtered on an Ultrafree-MC GV Centrifugal filter (Millipore, UFC30GV0S). Superose 6 Increase HR 10/300 column (Cytiva) was equilibrated with SEC buffer [25 mM HEPES (pH 7.5), 150 mM NaCl, 0.5 mM TCEP] at 0.4 ml/min before a total of 10 mg of protein from HEK293 cell lysate was applied to the column. Fractions of 0.5 mL were collected and analyzed by western blot.

### Generation of stable cell lines for BioID assay

Flp-In T-REx HEK293 cells were transfected using jetPRIME transfection reagent (Polyplus) according to manufacturer’s specifications. Briefly, cells were seeded in 6-well plates with DMEM supplemented with 5% FBS, 5% Cosmic calf serum and 100 U/mL Pen/Strep and transfected with 100 ng of pcDNA5-BAIT_PROTEIN-BirA*-FLAG and 1 μg of POG44 (Flp recombinase). Transfected cells were passaged into 10 cm plates and 48 h post-transfection treated with hygromycin B (200 µg/mL) for stable selection of integrated cells. Selection media was changed every 2-3 days until clear visible colonies were present. Mixed clonal populations were pooled and scaled up into three 15 cm plates (1 to freeze and 2 for BioID).

### BioID; Affinity purification (AP) and Trypsin digestion

For BioID experiments, stable cells were grown to ∼75% confluency. Bait expression vectors and biotin were induced simultaneously (1 µg/ml tetracycline, 50 µM biotin). After 24 h of treatment, cells were rinsed once on the plate with ∼20 ml PBS, then scraped into 1 ml of PBS. Cell pellets were collected by centrifugation (500 g for 5 min) and stored at –80°C for further processing. Cell pellets were thawed on ice and tared weight calculated. A 4:1 (v/w) ratio of ice-cold lysis buffer was added to the cells (50 mM Tris-HCl, pH 7.5, 150 mM NaCl, 1% NP40, 0.4% SDS, 1.5 mM MgCl_2_, 1 mM EGTA, benzonase, Sigma protease inhibitors). Cells were dispersed with a P1000 pipette tip (∼10-15 aspirations) and subjected to a rapid freeze/thaw cycle (dry ice to 37°C water bath). Lysates were rotated at 4°C for 30 min and then centrifuged at 16,000 g for 20 min at 4°C. Supernatant was collected (with 20 µL aliquot saved for western blot) into new tubes for affinity purification (AP). Samples were incubated with 20 µL (packed beads) of streptavidin-Sepharose (GE) (equilibrated in lysis buffer) with rotation overnight at 4°C. Beads were collected (500 g for 2 min), the supernatant discarded, and the beads transferred to new tubes in 500 µl of lysis buffer. Beads were washed once with SDS wash buffer (50 mM Tris-HCl, pH 7.5, 2% SDS), 2x with lysis buffer, and 3x with 50 mM ammonium bicarbonate, pH 8.0 (ABC) (all wash volumes = 500 µL with centrifugations at 500 g for 30 s). Beads were resuspended in 100 µl of ABC containing 1 µg of sequencing grade trypsin and gently mixed at 37°C for 4 h. 1 µg fresh trypsin was added, and the samples were rotated overnight. Supernatant was collected (500 g for 2 min) and the beads washed with 100 µl of molecular biology grade H_2_O and pooled with peptides. Digestion was terminated by acidification with formic acid (50 µL of 10% stock = 2% final concentration). Samples were then centrifuged (16,000 g for 5 min) and ∼90% of the sample was transferred to a new tube and dried with vacuum centrifugation.

### Mass spectrometry (MS)

Each sample (5 μL in 2 % formic acid; corresponding to 1/8th of a 15 cm tissue culture dish) was directly loaded at 800 nL/min onto an equilibrated HPLC column. The peptides were eluted from the column over a 90 min gradient generated by a Eksigent ekspert™ nanoLC 425 (Eksigent, Dublin CA) nano-pump and analysed on a TripleTOF 6600 instrument (SCIEX, Concord, Ontario, Canada). The gradient was delivered at 400 nL/min starting from 2% acetonitrile with 0.1% formic acid to 35% acetonitrile with 0.1% formic acid over 90 min followed by a 15 min clean-up at 80% acetonitrile with 0.1% formic acid, and a 15 min equilibration period back to 2% acetonitrile with 0.1% formic acid, for a total of 120 min. To minimize carryover between each sample, the analytical column was washed for 2 hours by running an alternating sawtooth gradient from 35% acetonitrile with 0.1% formic acid to 80% acetonitrile with 0.1% formic acid at a flow rate of 1500 nL/min, holding each gradient concentration for 5 min. Analytical column and instrument performance were verified after each sample by loading 30 fmol bovine serum albumin (BSA) tryptic peptide standard with 60 fmol α-casein tryptic digest and running a short 30 min gradient. TOF MS mass calibration was performed on BSA reference ions before running the next sample to adjust for mass drift and verify peak intensity. Samples were analyzed data dependent acquisition (DDA) mode. The DDA method consisted of one 250 milliseconds (ms) MS1 TOF survey scan from 400–1800 Da followed by ten 100 ms MS2 candidate ion scans from 100–1800 Da in high sensitivity mode. Only ions with a charge of 2+ to 5+ that exceeded a threshold of 300 cps were selected for MS2, and former precursors were excluded for 7 seconds after one occurrence.

### MS data analysis

All raw (WIFF and WIFF.SCAN) files were saved in the Gingras lab local interaction proteomics LIMS, ProHits (Liu et al., 2010). mzXML files were generated from raw files using the ProteoWizard converter converter (v3.0.4468) and SCIEX converter (v1.3 beta), implemented within ProHits. The searched database contained the human complement of the RefSeq protein database (version 57) complemented with SV40 large T-antigen sequence (72,226 sequences searched; including reversed sequences). mzXML files were searched by Mascot (v2.3.02) and Comet (v2016.01 rev. 2) using the following parameters: up to two missed trypsin cleavage sites, methionine oxidation and asparagine/glutamine deamidation as variable modifications. The fragment mass tolerance was 0.15 Da and the mass window for the precursor was ±40 ppm with charges of 2+ to 4+ (both monoisotopic mass). Search engine results were analyzed using the Trans-Proteomic Pipeline (TPP v4.6 OCCUPY rev 3) (Deutsch et al., 2010) via iProphet (Shteynberg et al., 2011). SAINTexpress (v3.6.3) (Teo et al., 2014) was used to score proximity interactions from DDA data. SAINTexpress calculates, for each prey protein identified by a given bait, the probability of a true proximity interaction relative to negative control runs using spectral counting as a proxy for abundance. Bait runs (two biological replicates each) were compared against eight negative control runs consisting of four BirA*-FLAG-only samples and four 3xFLAG-only samples, which were compressed to four “virtual controls” to maximize stringency of scoring. Preys with a false discovery rate (FDR) ≤ 1% (Bayesian estimation based on distribution of the Averaged SAINT scores across both biological replicates) were considered high-confidence proximity interactions.

## Acknowledgments

We thank Cassandra Wong for technical assistance. This work was supported by grants from the Canadian Institutes of Health Research (FDN-148423; to N.S.) and the Biotechnology and Biological Sciences Research Council (BB/W008165/1; to S.M.J.). J-H.C. is supported by the Basic Science Research Program through the National Research Foundation of Korea funded by the Ministry of Education (2020R1A6A3A03040141). J.L. is supported by funding from the China Scholarship Council. The graphical abstract was generated with BioRender.com.

## Author contributions

J-H.C. and J.L. performed most of the experiments and data analyses. X.Z. performed the virus-related experiments. G.G.H. performed the BioID assay. A.P. performed the size exclusion chromatography. R.L., S.G., and Y.A. assisted with experiments. J-H.C., J.L., X.Z., A.C.G., T.A., T.M.S., T.D., N.S., and S.M.J. contributed to the design and analysis of the experiments. J-H.C., J.L., X.Z., N.S., and S.M.J. contributed to the conception of the study. N.S. and S.M.J. supervised the project. J-H.C., J.L., X.Z., and

S.M.J. wrote the manuscript. All authors edited and approved the manuscript.

## Supplementary Figures Legends

**Supplementary Figure 1; related to** **Figure 1**. Alignment of the human GIGYF1 (NP_001362694.1) and GIGYF2 (NP_001096616.1) protein sequences by multAlin software (Corpet, 1988).

**Supplementary Figure 2. Translational repression of target mRNAs by GIGYF1; related to** **Figure 1**. (**A**) RT-qPCR analyses of samples described in Figure 1D. *GAPDH* mRNA level was used as a control for normalization. Data are presented as mean ± SD (n=3). ns = non-significant, analyzed by one-way ANOVA with Bonferroni’s post-hoc test. (**B**) Western blot analysis with the indicated antibodies of WT, 4EHP-KO, GIGYF1-KO, and GIGYF2-KO A549 cell lysates. (**C**) ELISA measurement of IFN-β production in WT, 4EHP-KO, GIGYF1-KO, and GIGYF2-KO A549 cells following 6 h of treatment with 1 μg/mL poly(I:C). Data are presented as mean ± SD (n=3). **P < 0.01; one-way ANOVA with Bonferroni’s post-hoc test. (**D**) Polysome profiling using WT and GIGYF1-KO HEK293 cells with or without poly(I:C) treatment. 1 μg/mL HMW poly(I:C) was used to stimulate the cells for 6 h. (**E**) Polysome (fractions 6-12) / monosome (fractions 3-5) (P/M) ratio of samples described in D. Data are presented as mean ± SD. ns = non-significant, **P < 0.01; two-way ANOVA with Bonferroni’s post-hoc test. (**F**) Co-immunoprecipitation assay for detection of interactions between GIGYF1 and 4EHP. HA-4EHP and v5-GIGYF1 or the 4EHP-binding mutant v5-GIGYF1 (v5-GIGYF1 Mut) were co-expressed in HEK293 cells. Cell lysates were immunoprecipitated with anti-v5 antibody. Immunoblotting was performed with the indicated antibodies. Inputs represent 2% of the total lysate used in the IP assay. (**G**) Western blot analysis with the indicated antibodies of cell lysates described in Fig. 1J.

**Supplementary Figure 3. Enhanced translational repression of target mRNAs by GIGYF1, compared with GIGYF2; related to** **Figure 2**. (**A**) WT, 4EHP-KO, GIGYF1-KO, and GIGYF2-KO HEK293 cells were transfected with psiCHECK2-RL-*Ifnb1* 3’ UTR reporter or the psiCHECK2 reporter (as control). RL and FL activities were measured 24 h after transfection. The RL/FL ratio in psiCHECK2-RL-*Ifnb1* 3’ UTR reporter–expressing cells was normalized to the psiCHECK2-expressing cells. Data are presented as mean ± SD (n=3). ns = non-significant, ****P < 0.0001; two-way ANOVA with Bonferroni’s post-hoc test. (**B**) Western blot analysis of cell lysate from Figure 2C with the indicated antibodies. (**C** & **D**) Tether-function assays for measurement of repression of the deadenylation-permissive RL-5boxB-polyA (**C**) or deadenylation-resistant RL-5BoxB-HhR reporters (**D**) upon coexpression with increasing amounts of λN-v5-GIGYF1 or λN-v5-GIGYF2 plasmids in HEK293 cells. Data are presented as mean ± SD (n=2). *P < 0.05, ****P < 0.0001; two-way ANOVA with Bonferroni’s post-hoc test. (**E-G**) Western blotting with the indicated antibodies using lysates from cells shown in Figure 2E, F, and H, respectively.

**Supplementary Figure 4. Interactions between GIGYF1 and eIF3 subunits; related to** **Figure 3**. (**A**) *Left*: PLA for detection of GIGYF1-eIF3G interaction in HEK293T cells transfected with vectors expressing v5-GIGYF1 or v5-GIGYF2 along with FLAG-eIF3G. 24 h post-transfection cells were fixed and subjected to PLA using FLAG and v5 antibodies. Scale bar = 10 µm. *Right*: The bar graphs represent the number of PLA signals from at least 30 cells, counted in each sample. n=5 independent experiments. Data are presented as mean ± S.E.M. ns= non-significant, ****P< 0.0001; unpaired t-test. (**B**) Western blot analysis of the samples in Figure 3B with the indicated antibodies. (**C**) Western blot analysis of the samples in Figure 3C with the indicated antibodies. (**D**) Western blot analysis of the samples in Supp. Fig. 4A with the indicated antibodies. (**E**) Fractionation of endogenous GIGYF1, GIGYF2, and the indicated subunits of eIF3 complex by size-exclusion chromatography. A total of 10 mg of proteins from HEK293 cells was loaded onto a Superose 6 column and ran at a flow rate of 0.4 mL/min. Fractions of 0.5 mL were collected, and 50 μL of each fraction was subjected to western blot analysis. The elution position of the molecular size markers is shown. (**F**) Schematic of the domain structures of full-length (FL) GIGYF1 and truncated isoforms A-E used in Figure 3D. (**G**) Western blot analysis of the samples in Figure 3D with the indicated antibodies. (**H**) Western blot analysis of the samples in Figure 3E with the indicated antibodies.

**Supplementary Figure 5. The C-terminal region of GIGYF1 mediates its interactions with eIF3D; related to** **Figure 3**. (**A**) *Left*: HEK293T cells were co-transfected with FLAG-eIF3D and the full-length or indicated GIGYF1 truncation mutants (see Supp. Fig. 4F for more details) or full-length GIGYF1 (as control). 24 h post-transfection cells were fixed and subjected to PLA using FLAG and v5 antibodies. PLA signals are shown in yellow. The nucleus and actin cytoskeleton were counterstained with DAPI and phalloidin (red), respectively. Scale bar = 10 µm. *Right*: The bar graphs represent the number of PLA signals from at least 20 cells, counted in each sample. n=3 independent experiments. ns = non-significant, ****P< 0.0001; one-way ANOVA with Bonferroni’s post-hoc test. (**B**) Western blot analysis of cell lysates from (A) with the indicated antibodies. (**C**) *Left*: PLA assay for detection of the interactions between eIF3D and the indicated two chimeric constructs described in Figure 2G. *Right*: The bar graphs represent the number of PLA signals from at least 20 cells, counted in each sample. n=3 independent experiments. Data are presented as mean ± S.E.M. ns= non-significant, ****P < 0.0001; one-way ANOVA with Bonferroni’s post-hoc test. (**D**) Western blot analysis of cell lysates from (C) with the indicated antibodies.

**Supplementary Figure 6. Ectopic expression of GIGYF1 reduces interactions of endogenous eIF3 and eIF4G1 proteins; related to** **Figure 4**. (**A**) Co-IP assay for detection of interactions of FLAG-eIF3D with eIF2α and TTP in GIGYF1-KO cells expressing v5-GIGYF1 or v5-Empty control plasmids. Whole-cell lysates were prepared 24 h after transfection and subjected to immunoprecipitation using FLAG antibody followed by blotting with the indicated antibodies. (**B**) Co-IP assay for detection of interactions of eIF3D with eIF3E and eIF4G1 in GIGYF1-KO cells expressing increasing amounts of GIGYF1. Whole-cell lysates were prepared 24 h after transfection and subjected to immunoprecipitation using anti-eIF3D antibody followed by blotting with the indicated antibodies. (**C**) Co-IP assay for detection of interactions of eIF3E with eIF3D and eIF4G1 in GIGYF1-KO cells expressing v5-Empty control or v5-GIGYF1. (**D**) Western blot analysis of expression of the indicated protein in samples from Figure 4C. (**E**) Western blot analysis of expression of the indicated protein in samples from Figure 4D. (**F**) Streptavidin-biotin RNA affinity purification assay with biotinylated *Ifnb1* 3’ UTR. Biotinylated *Ifnb1* 3’ UTR was incubated with GIGYF1-KO cell lysates in the presence or absence of 10X non-biotinylated *Ifnb1* 3’ UTR for 16 h at 4°C. The pulled-down proteins were subjected to western blotting and probed with the indicated antibodies. (**G**) Western blot analysis of cell lysates from Figure 4H. (**H**) *Left*: PLA assay for detection of interactions between v5-GIGYF1 and FLAG-eIF3G upon Poly(I:C) stimulation. PLA signals are shown in yellow, the nucleus and Actin cytoskeleton were counterstained with DAPI and phalloidin (red), respectively. Scale bar = 10 µm. *Right*: The bar graphs represent the number of PLA signals from at least 20 cells, counted in each sample. N=3 independent experiments. Data are presented as mean ± S.E.M. ns= non-significant, ****P < 0.0001; unpaired t-test. (**I**) Western blot analysis of cell lysates from (H). (**J**) Western blot analysis of cell lysates from Figure 4I.

**Supplementary Figure 7. Uncropped images of blots used in** **Figure 1** **to** **Figure 4** **and Supplementary Figures**.

**Supplementary Table 1. Combined data from two independent BioID experiments using N– or C-terminal tagged BirA*-GIGYF1 in HEK293 cells.** N and C (N– and C– terminal, respectively) indicate the location of BirA* fusion protein in relation to the bait protein.

**Supplementary Table 2. List of primers and sgRNAs used in this study**.

