## Supplementary Figures for "Repression of mRNA translation initiation by GIGYF1 via blocking the eIF3-eIF4G1 interaction"

#### Supplementary Figure. 1

Aligned sequences : GIGYF1, GIGYF2

- Identity : 580/1347 (43.1%)
- Similarity : 739/1347 (54.9%)

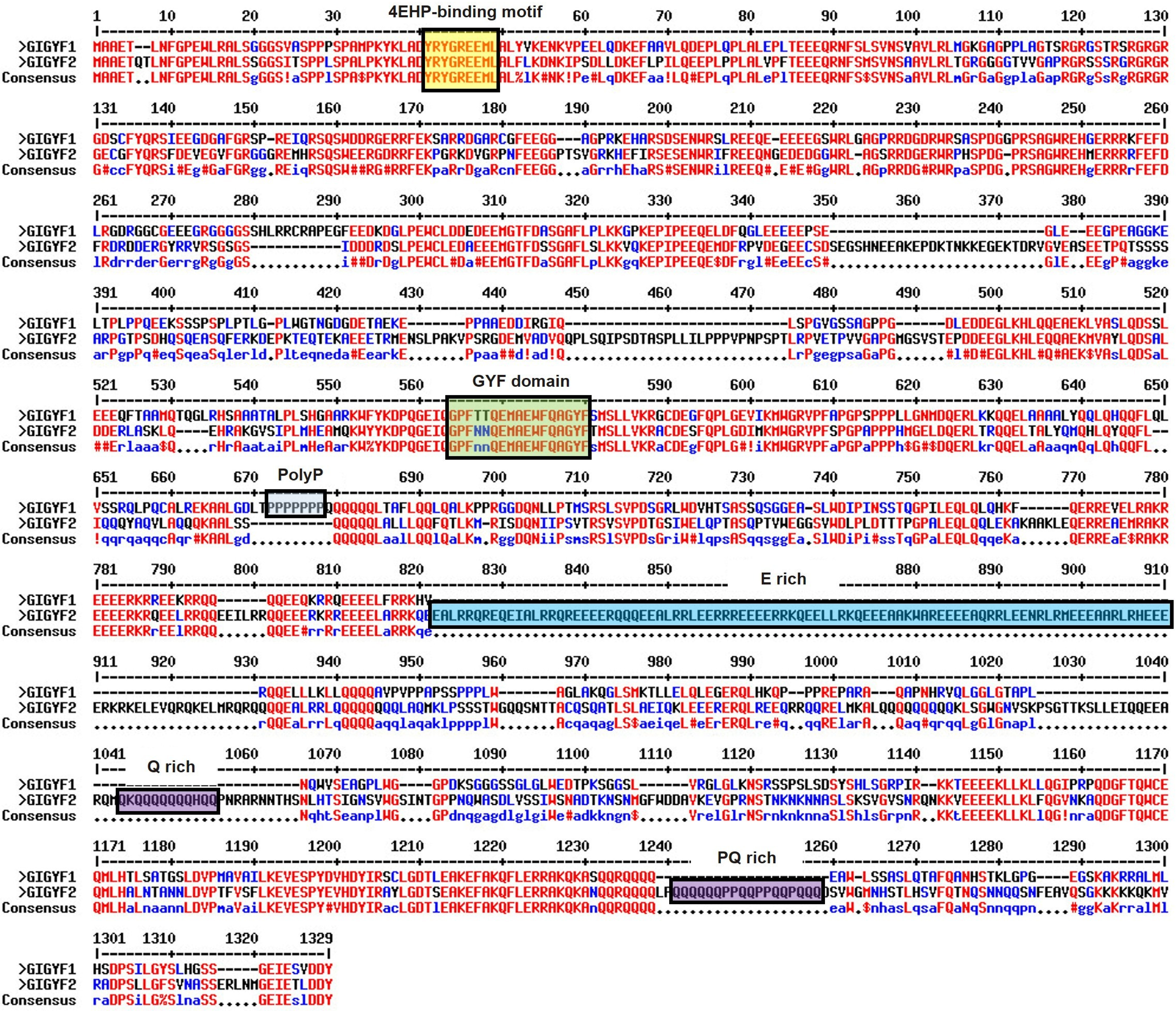

#### Supplementary Figure. 2

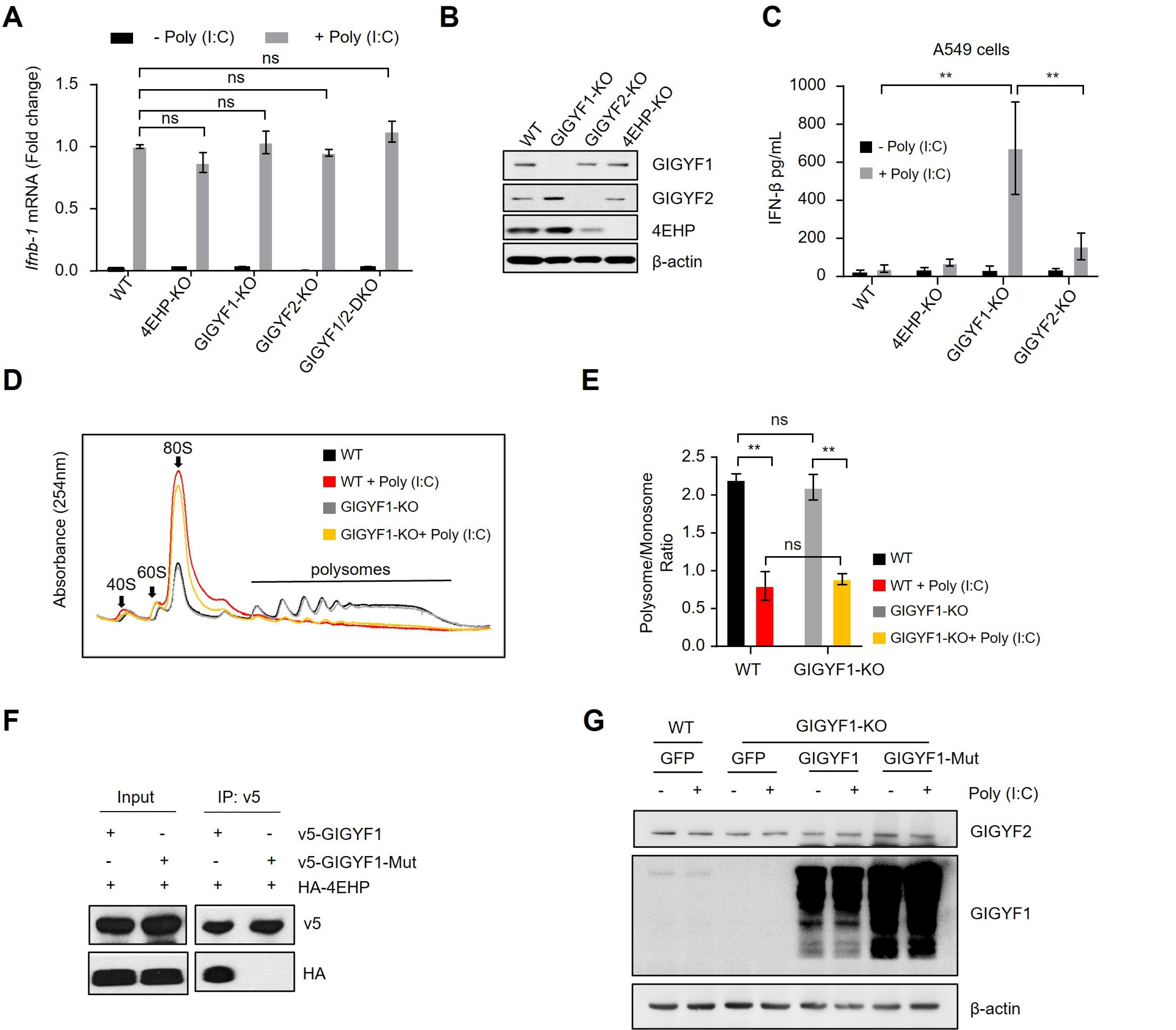

### Supplementary Figure. 3

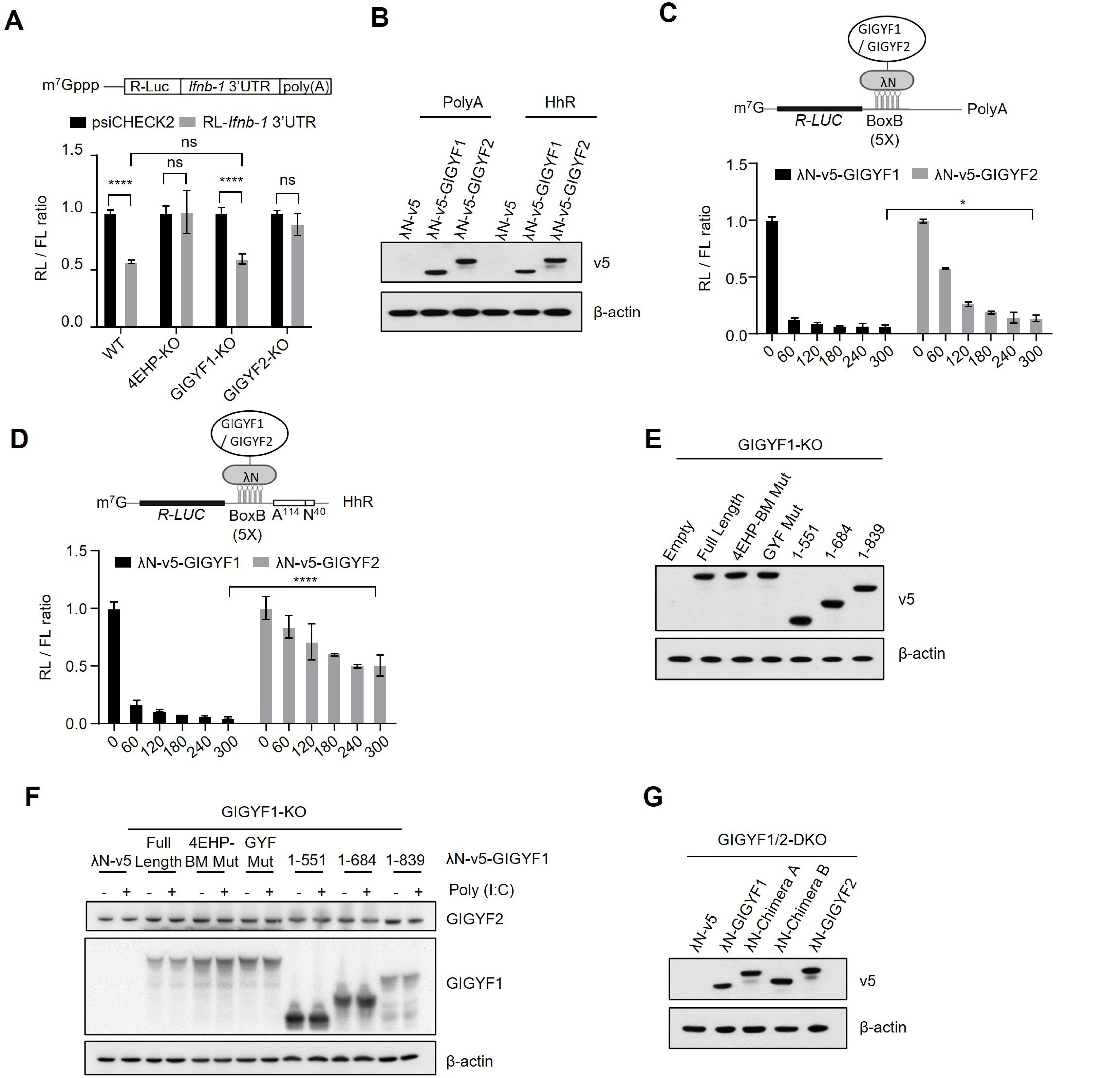

Supplementary Figure. 4

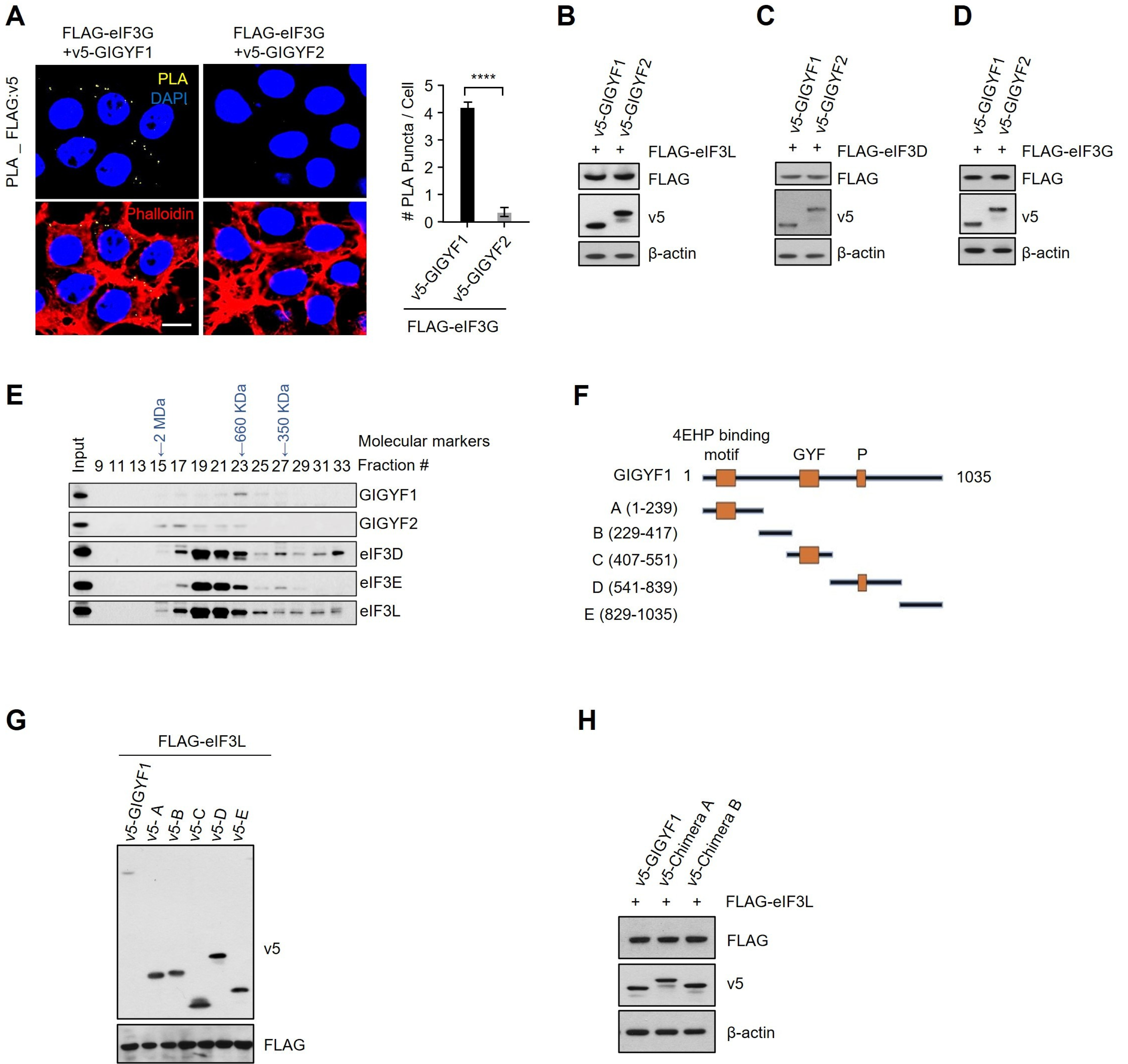

### Supplementary Figure. 5

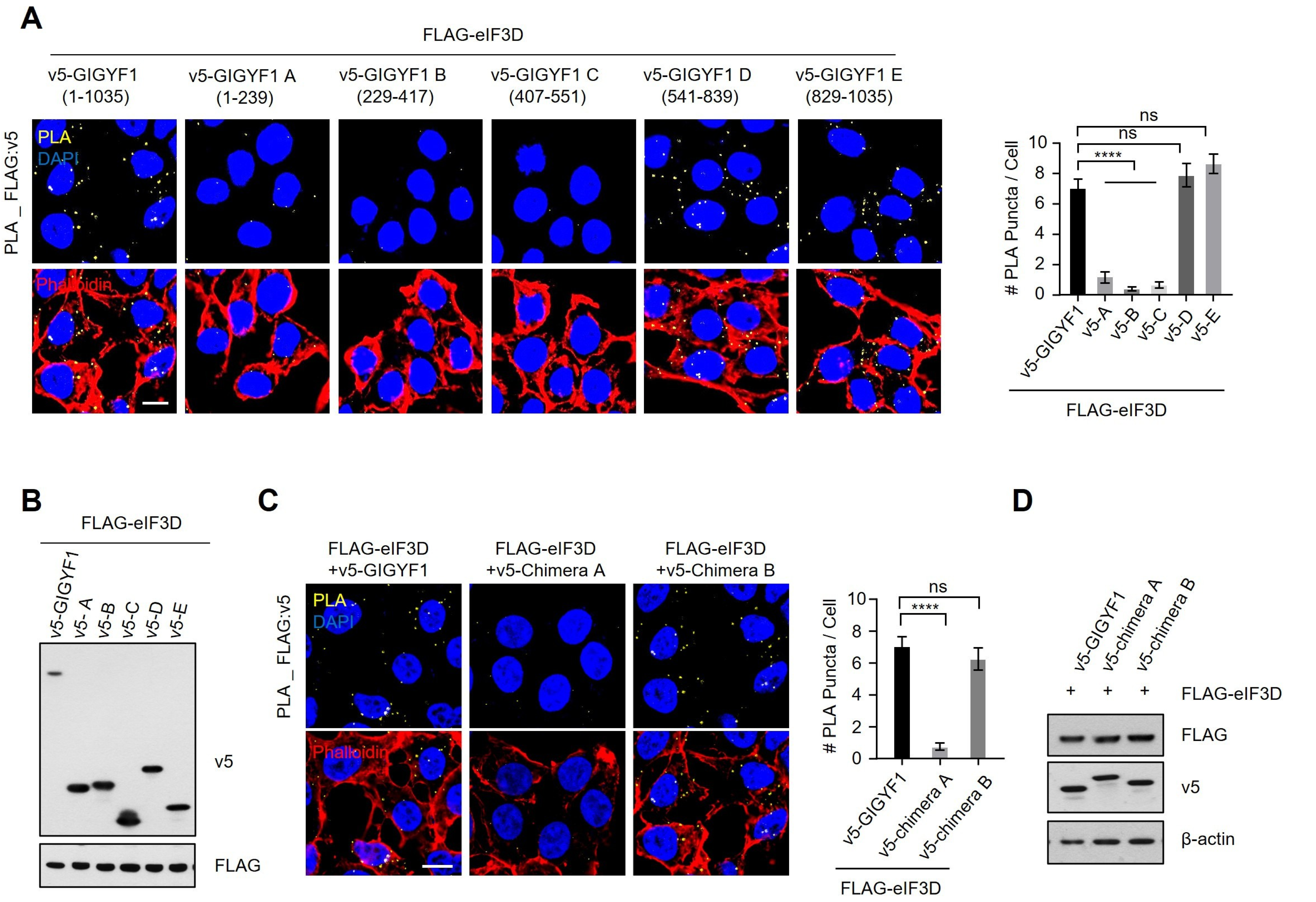

### Supplementary Figure. 6

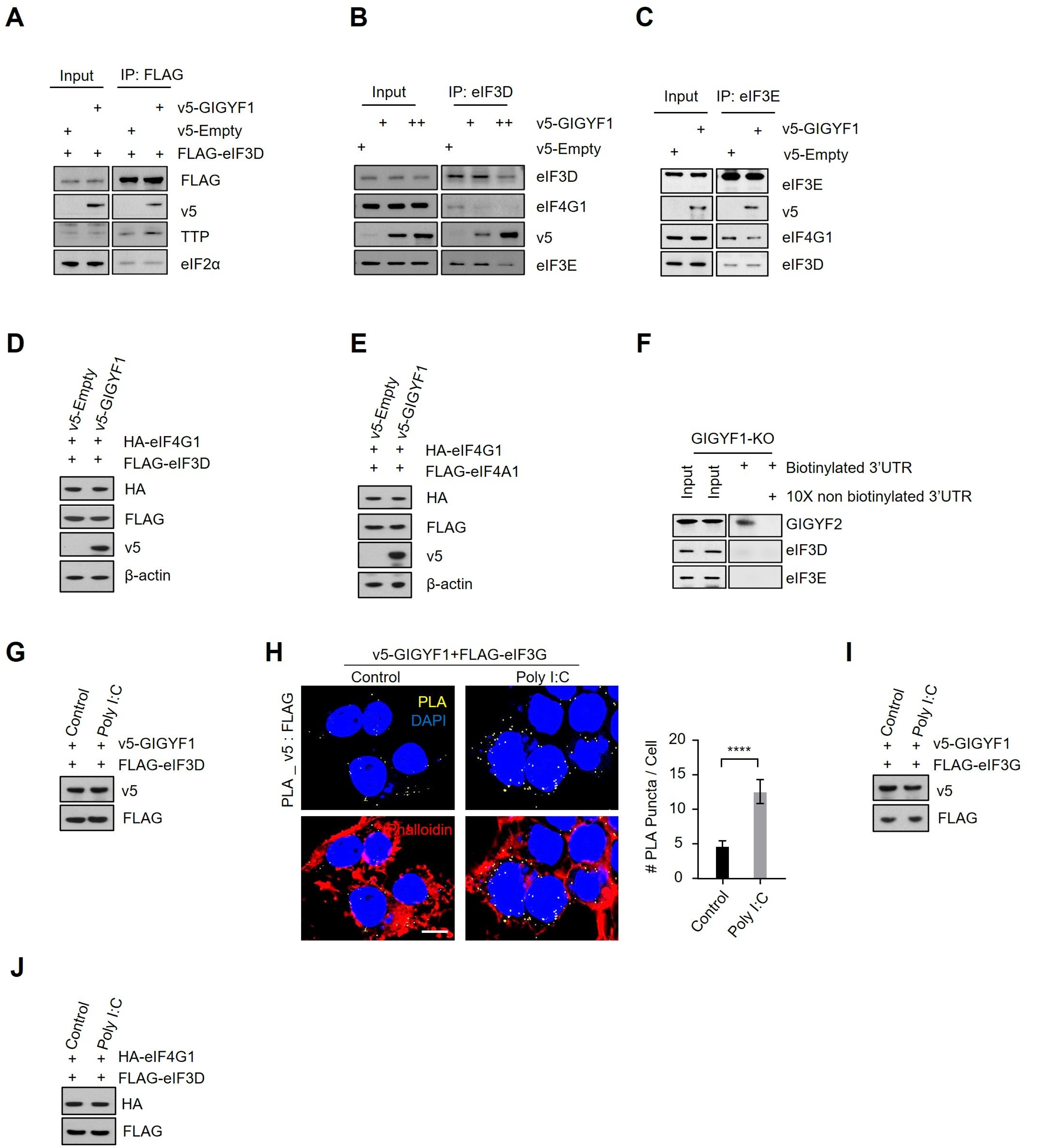

### Supplementary Figure. 7 ; Original blots for Figure. 1

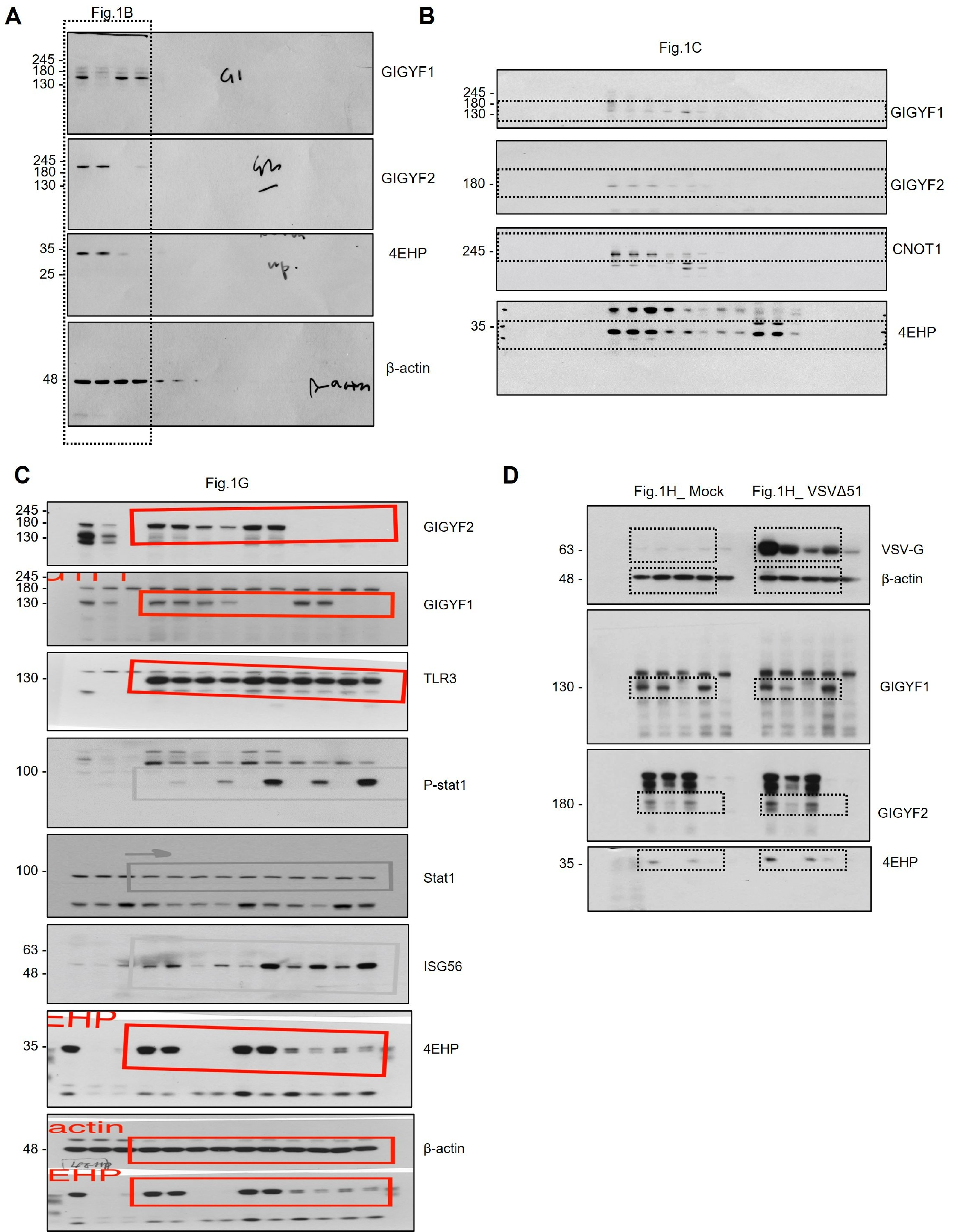

### Supplementary Figure. 7 ; Original blots for Supplementary Figure. 2

**A**

Fig.S2B

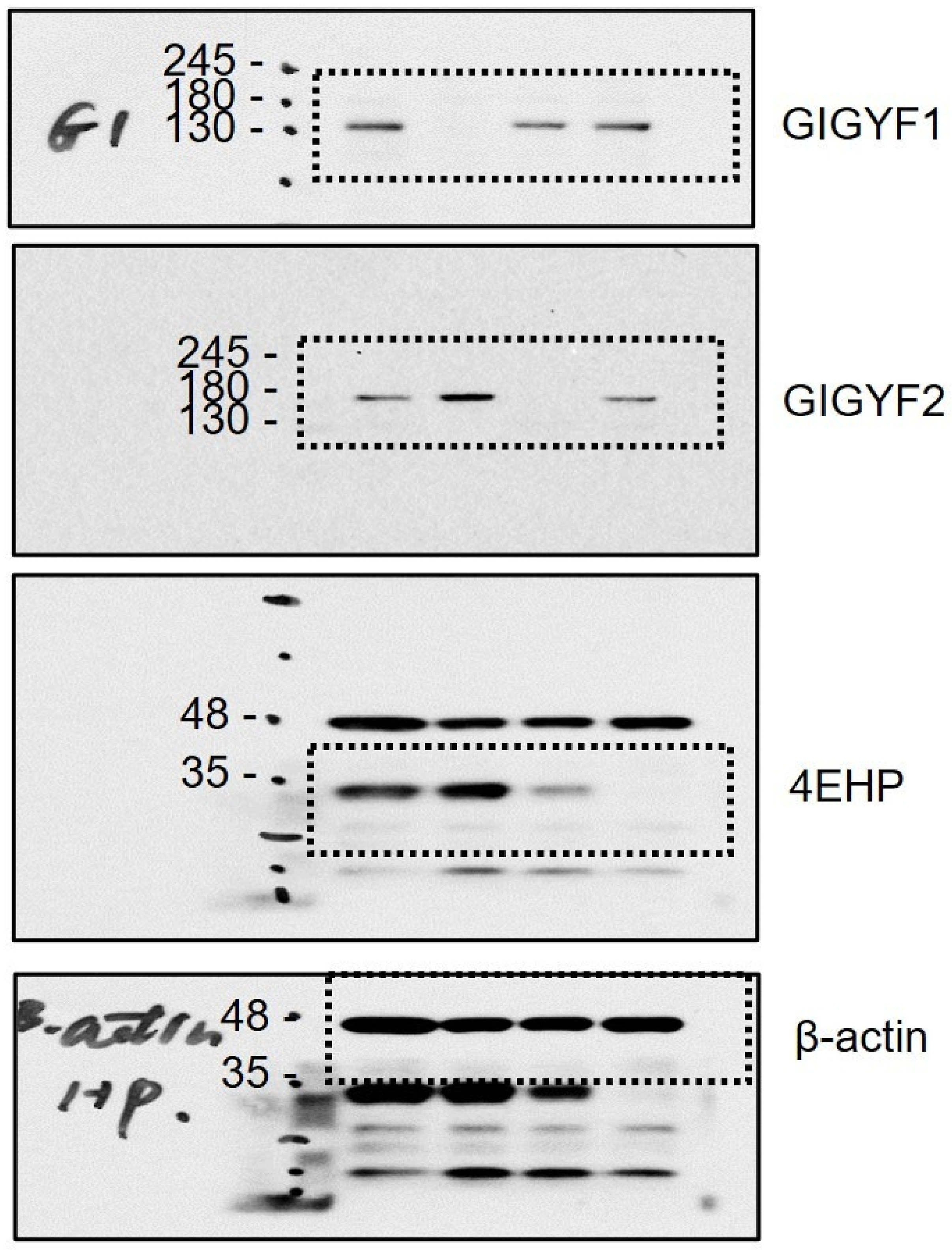


**B**

Fig.S2F_ Input

Fig.S2F_ IP : v5

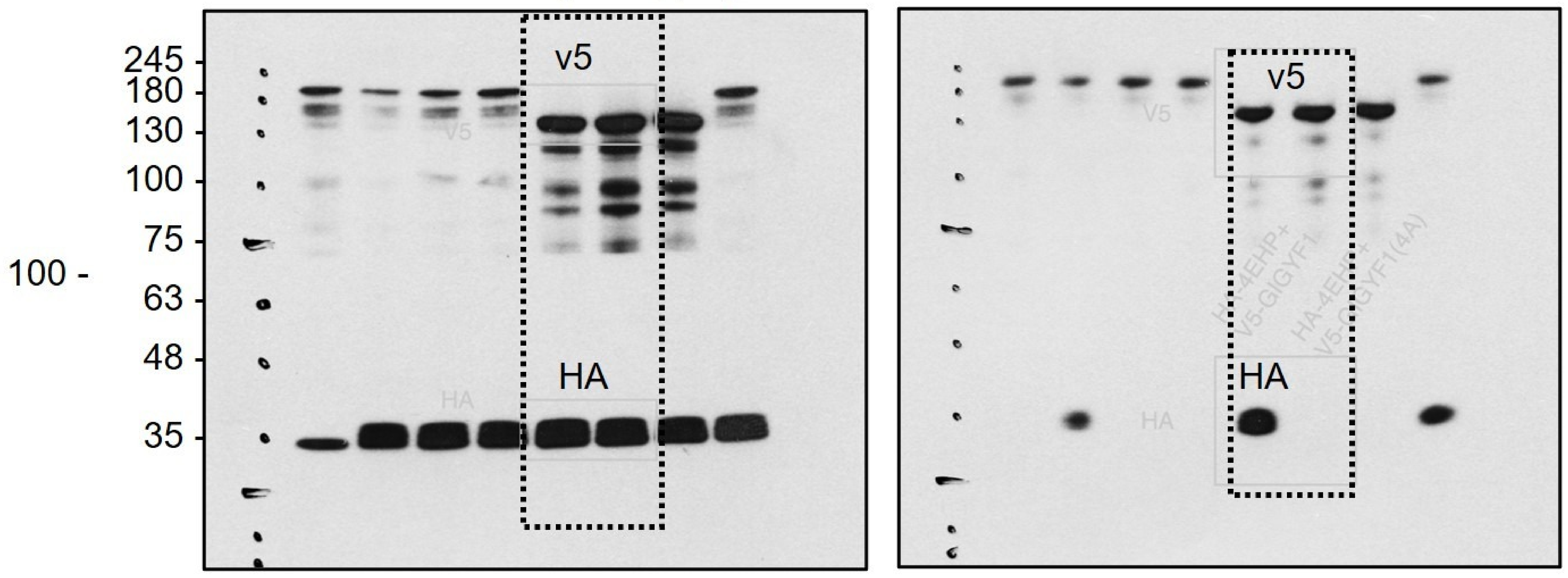


**C**

Fig.S2G

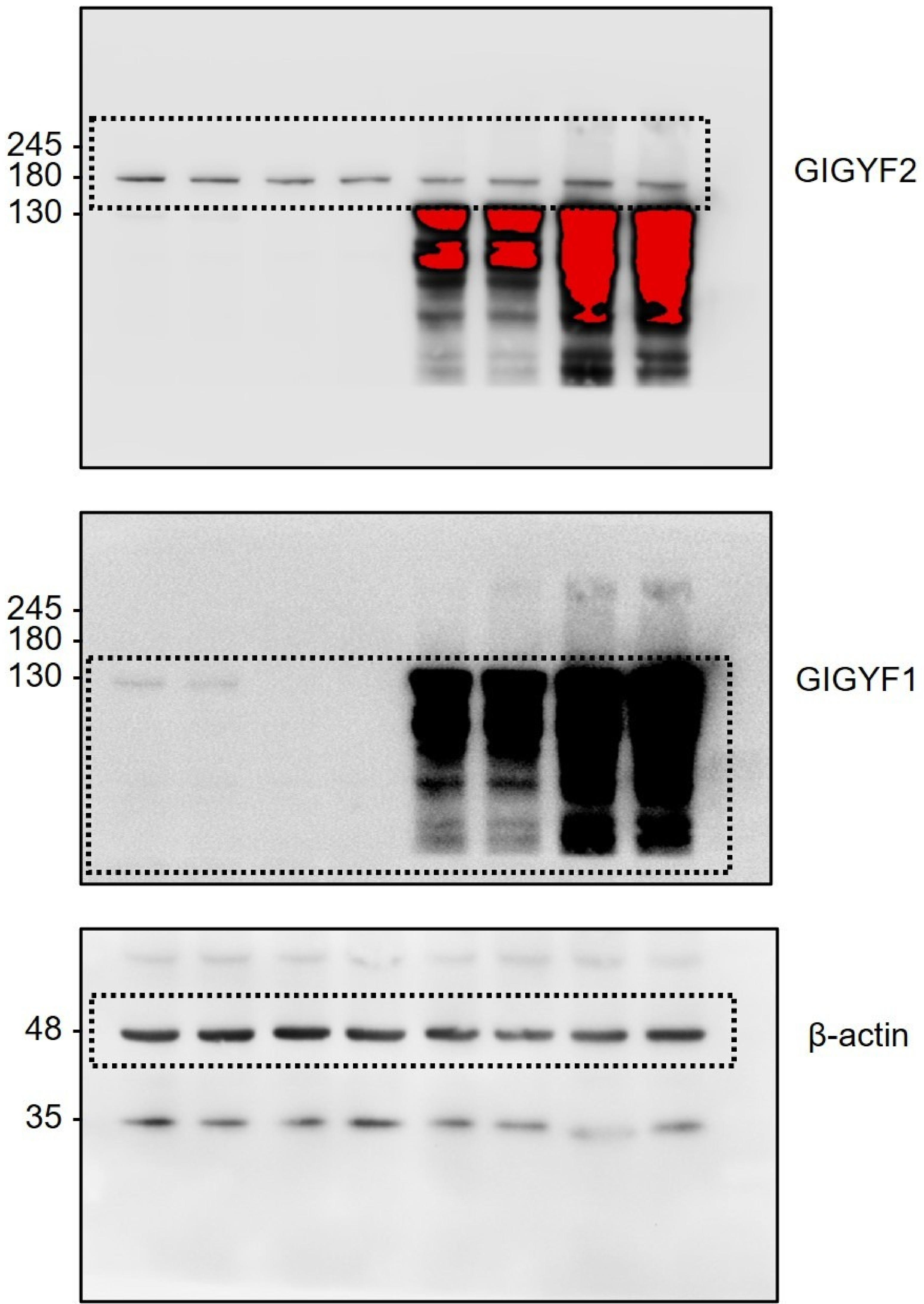

### Supplementary Figure. 7 ; Original blots for Supplementary Figure. 3

**A**

Fig.S3B

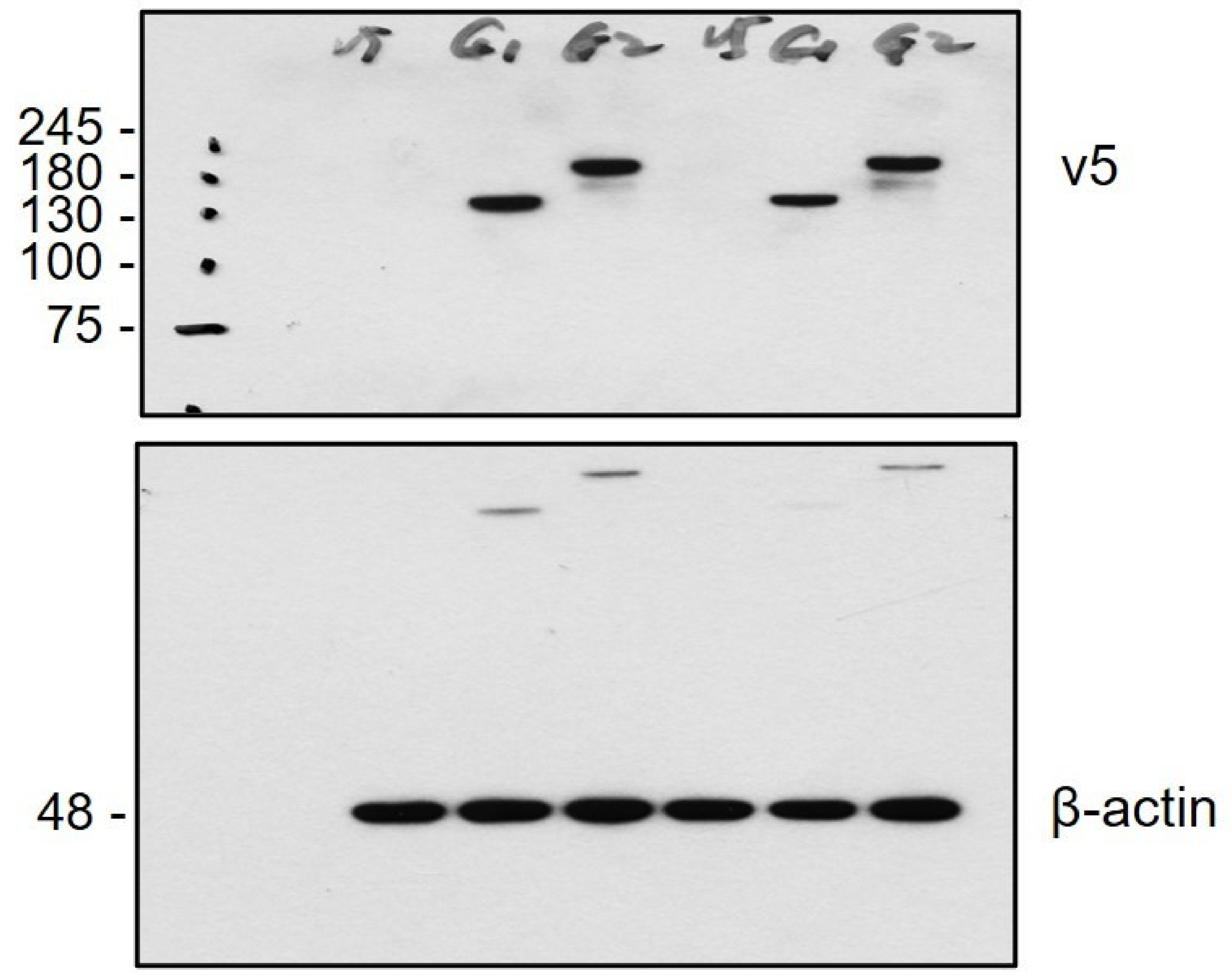


**B**

Fig.S3E

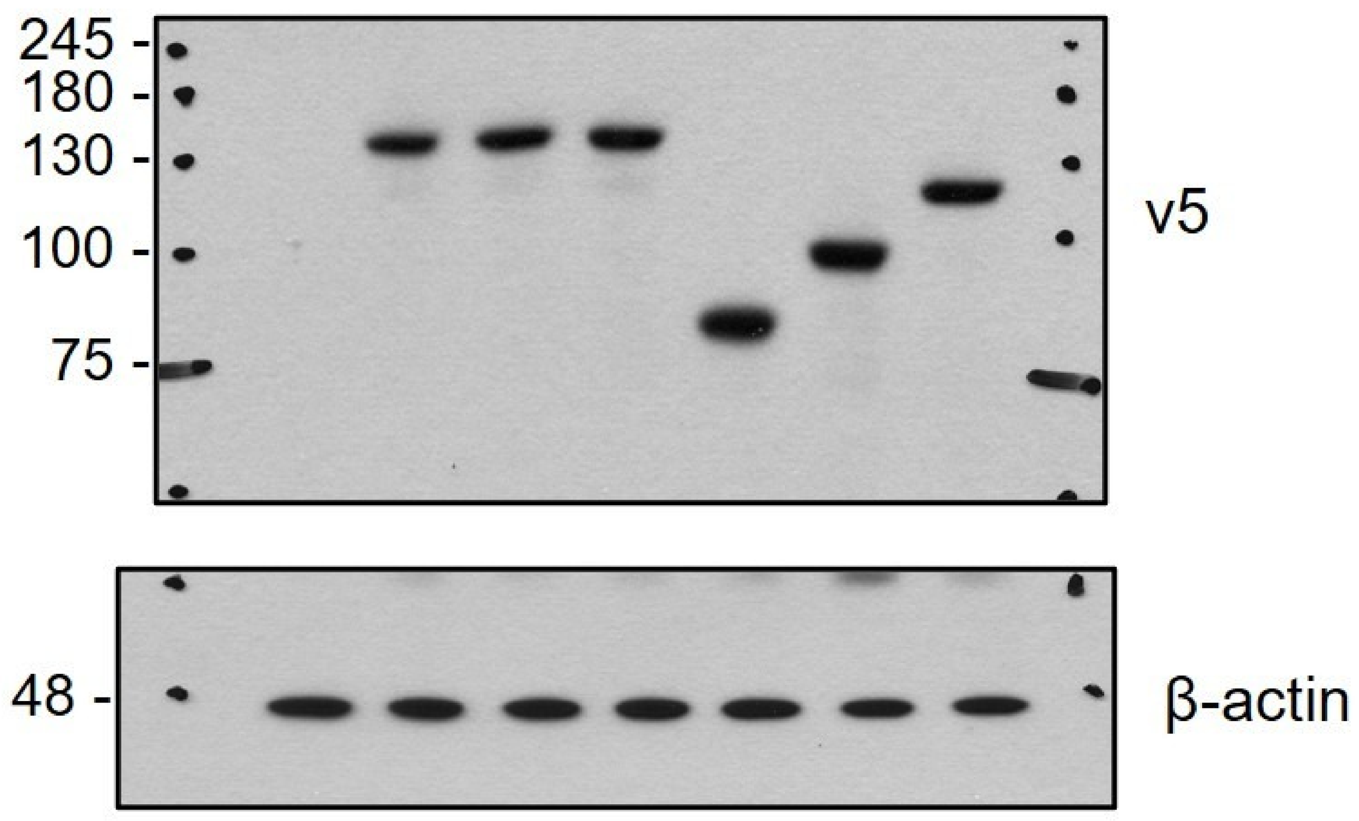


**C**

Fig.S3F

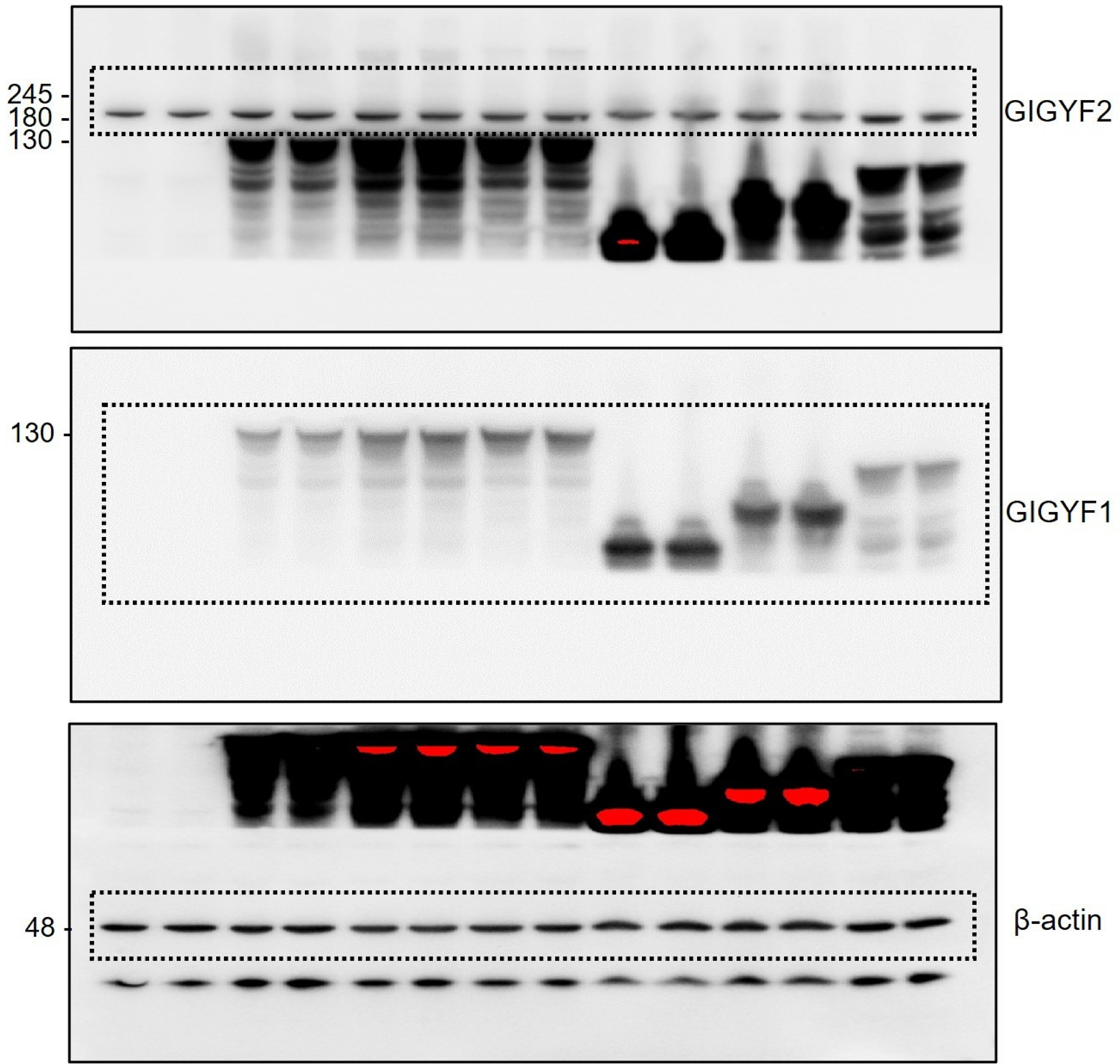


**D**

Fig.S3G

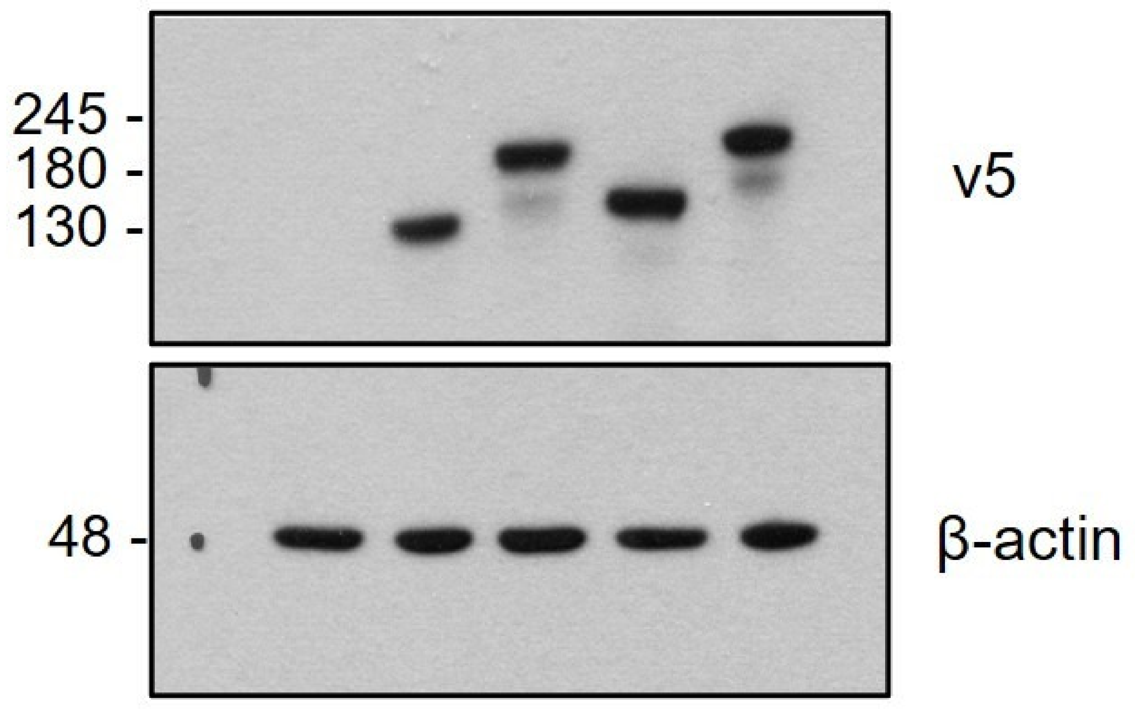

### Supplementary Figure. 7 ; Original blots for Figure. 3

## A

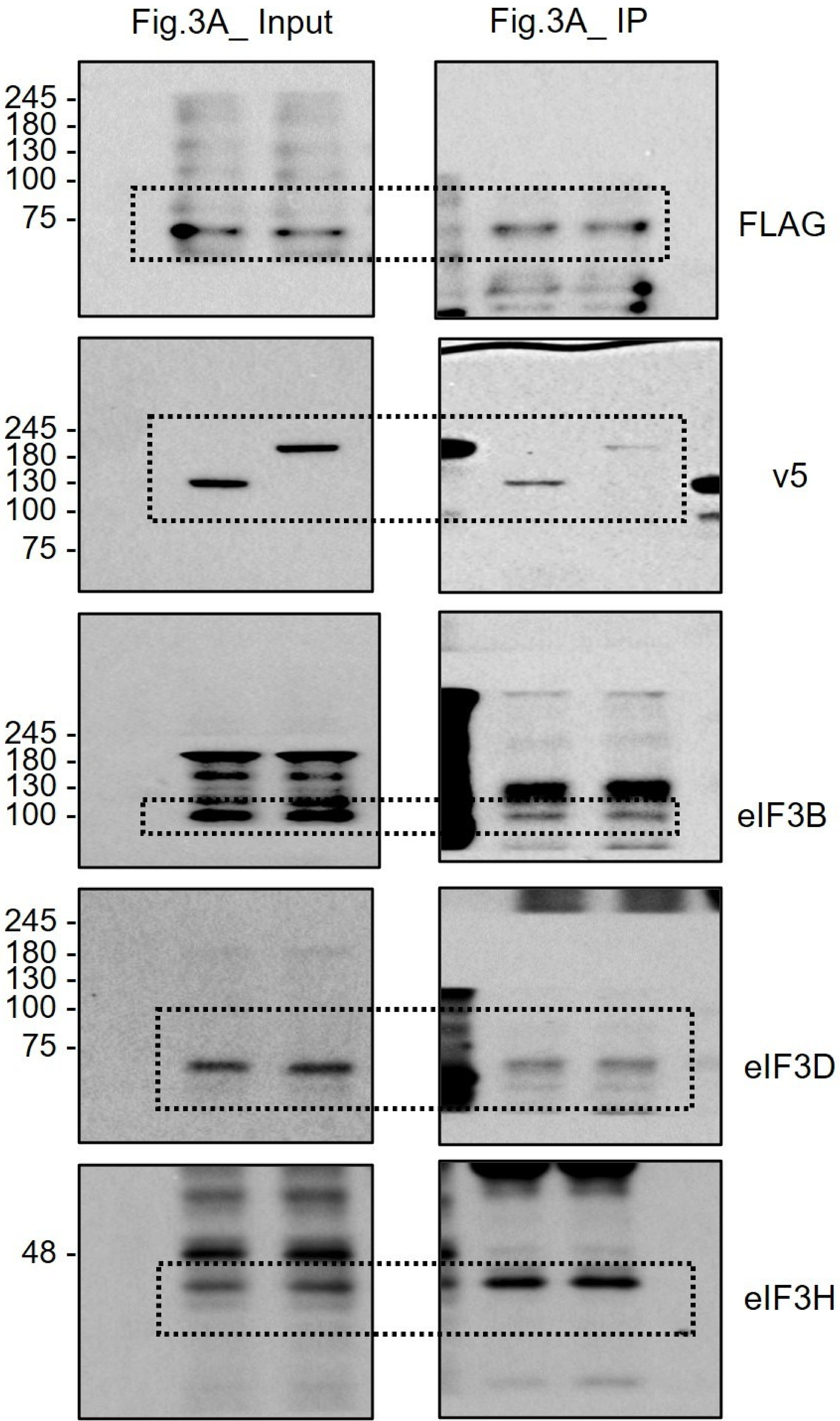

**Supplementary Figure. 7 ; Original blots for Supplementary Figure. 4**

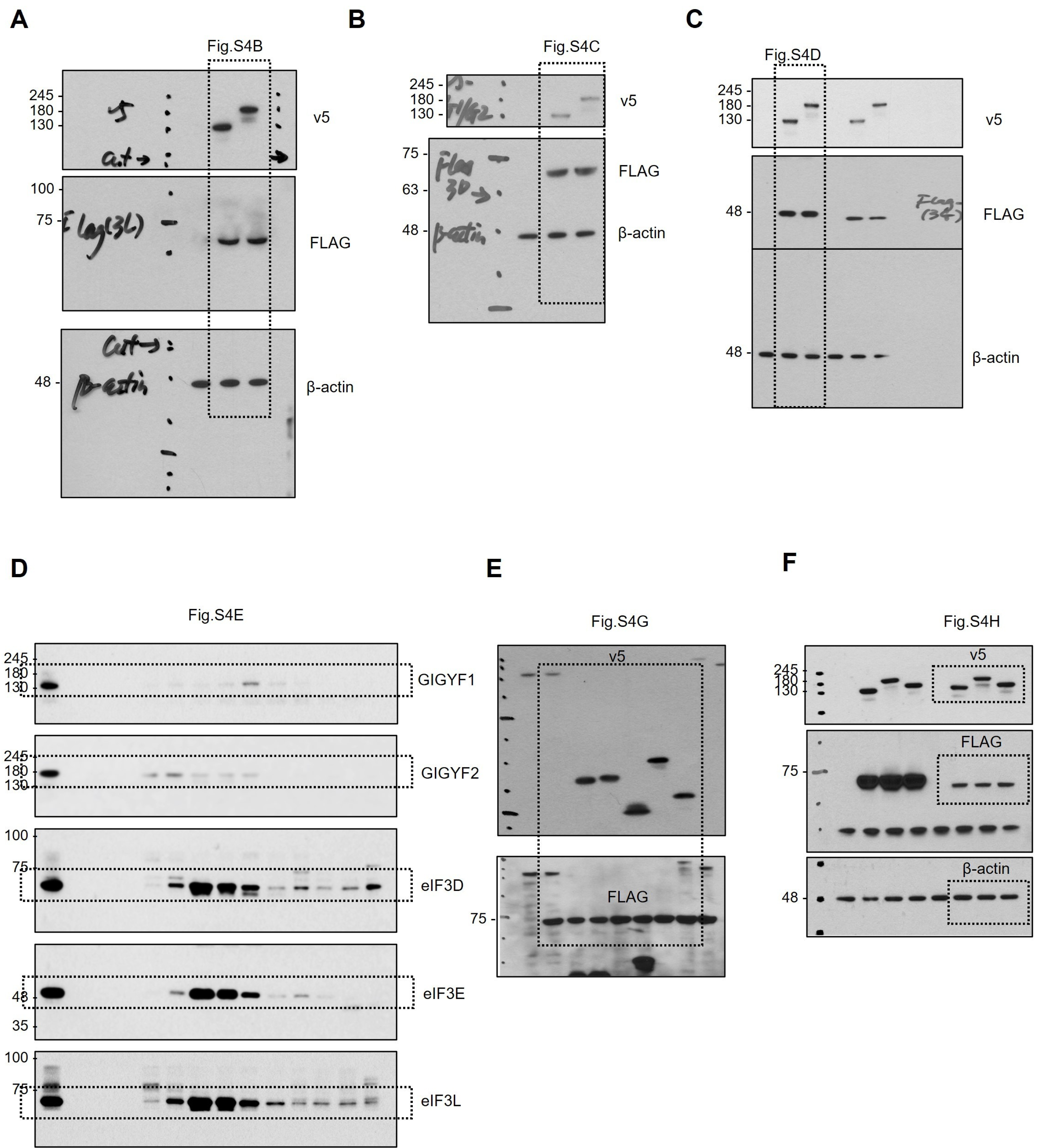

### Supplementary Figure. 7 ; Original blots for Supplementary Figure. 5

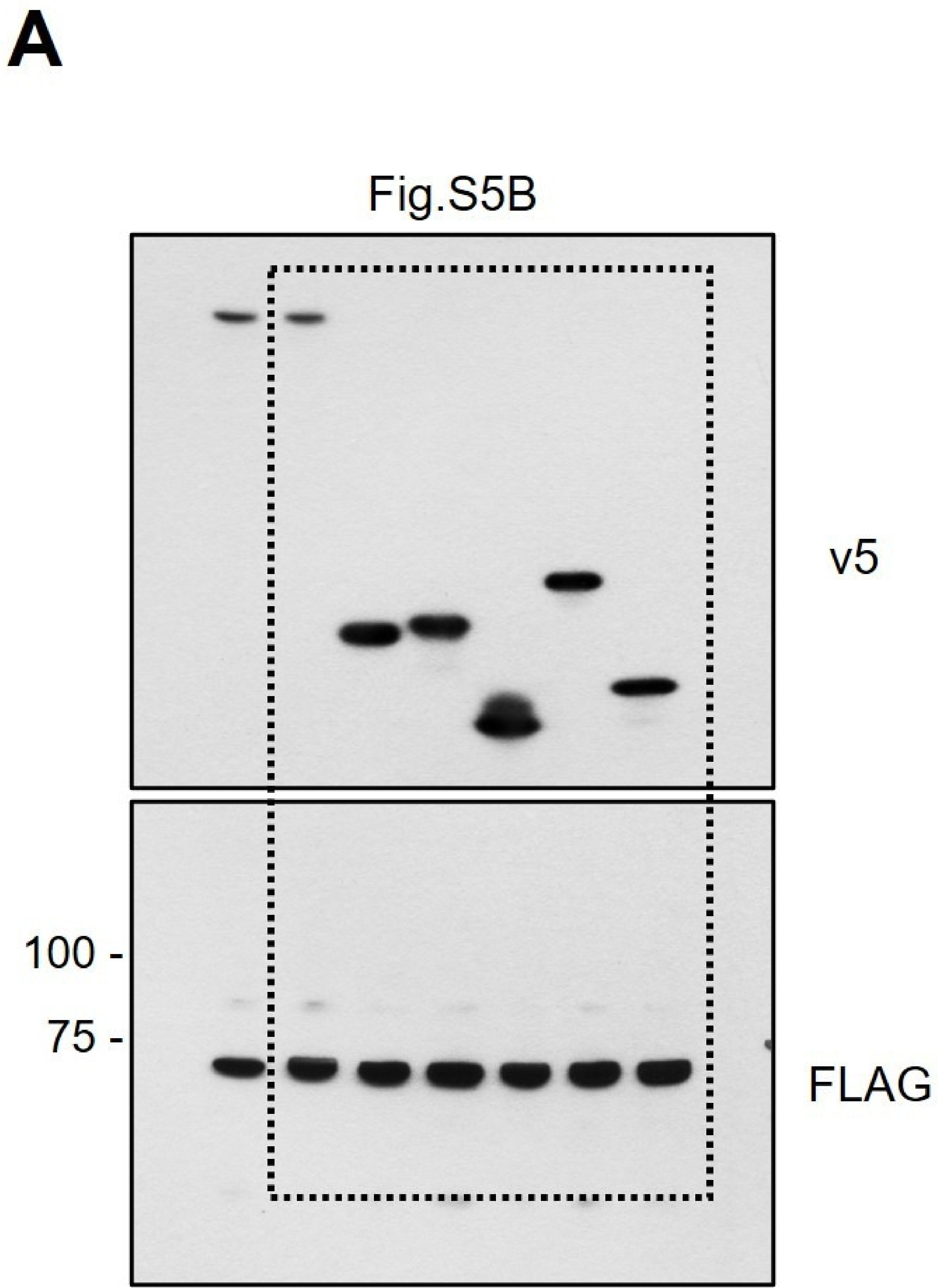


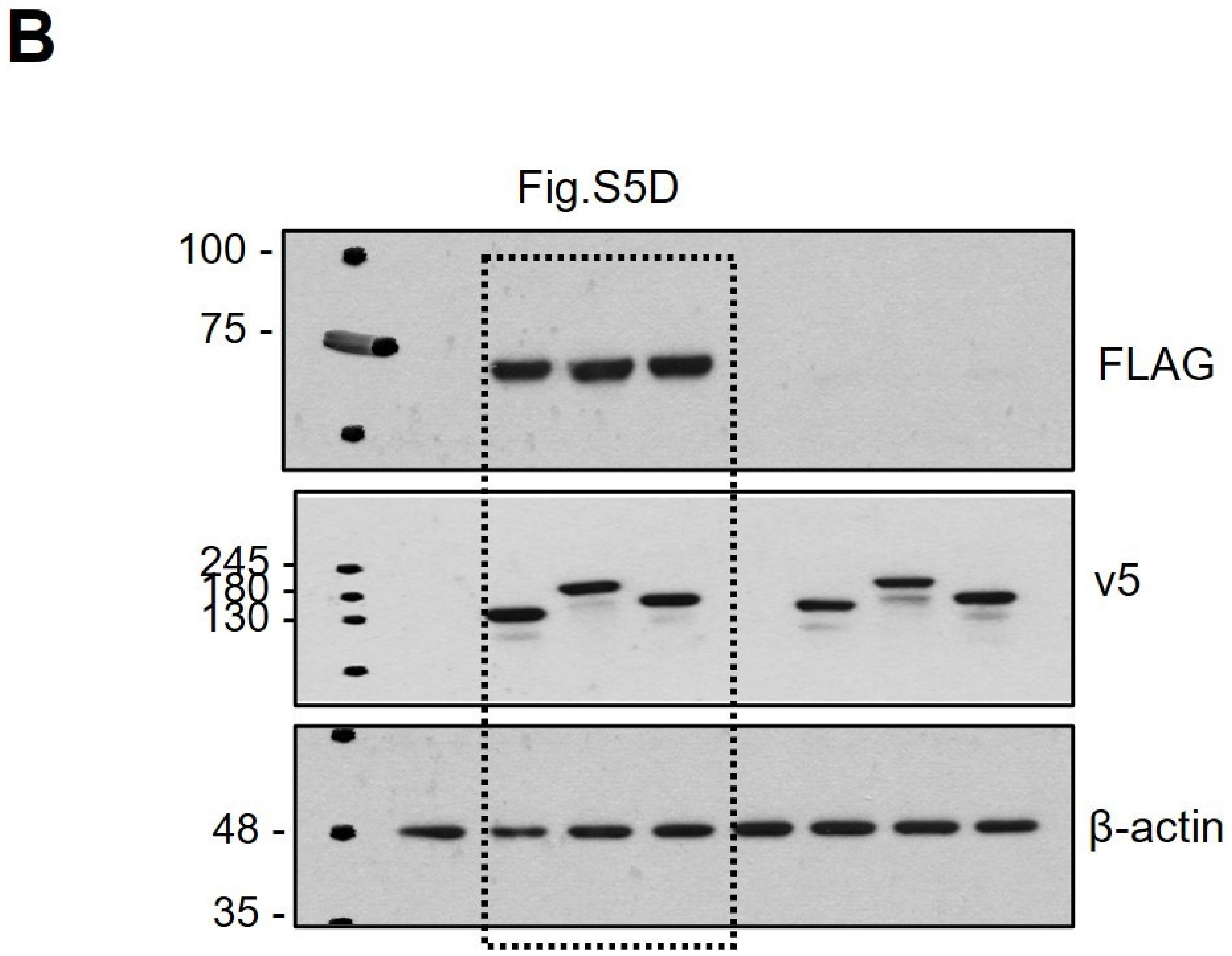

### Supplementary Figure. 7 ; Original blots for Figure. 4

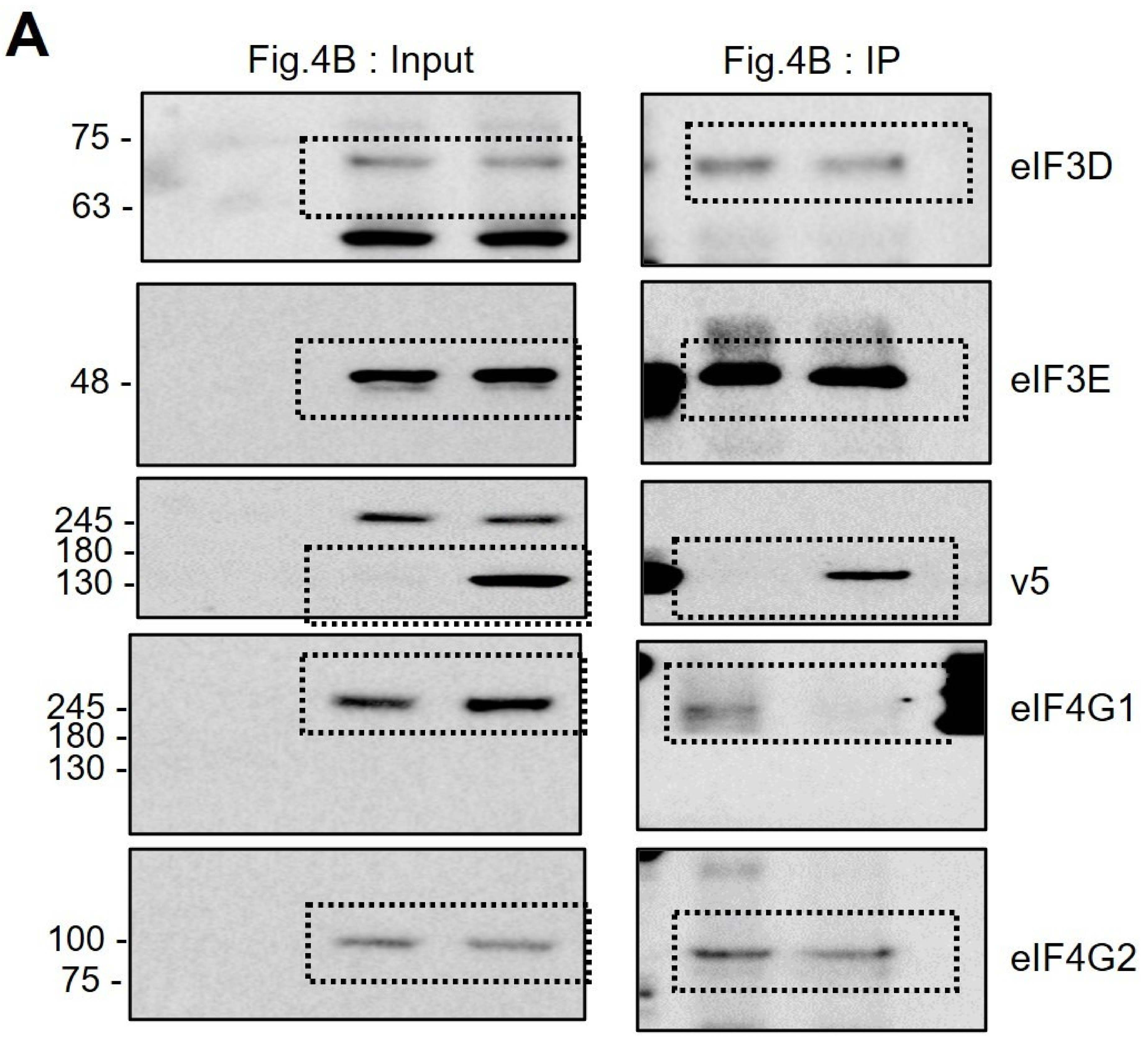


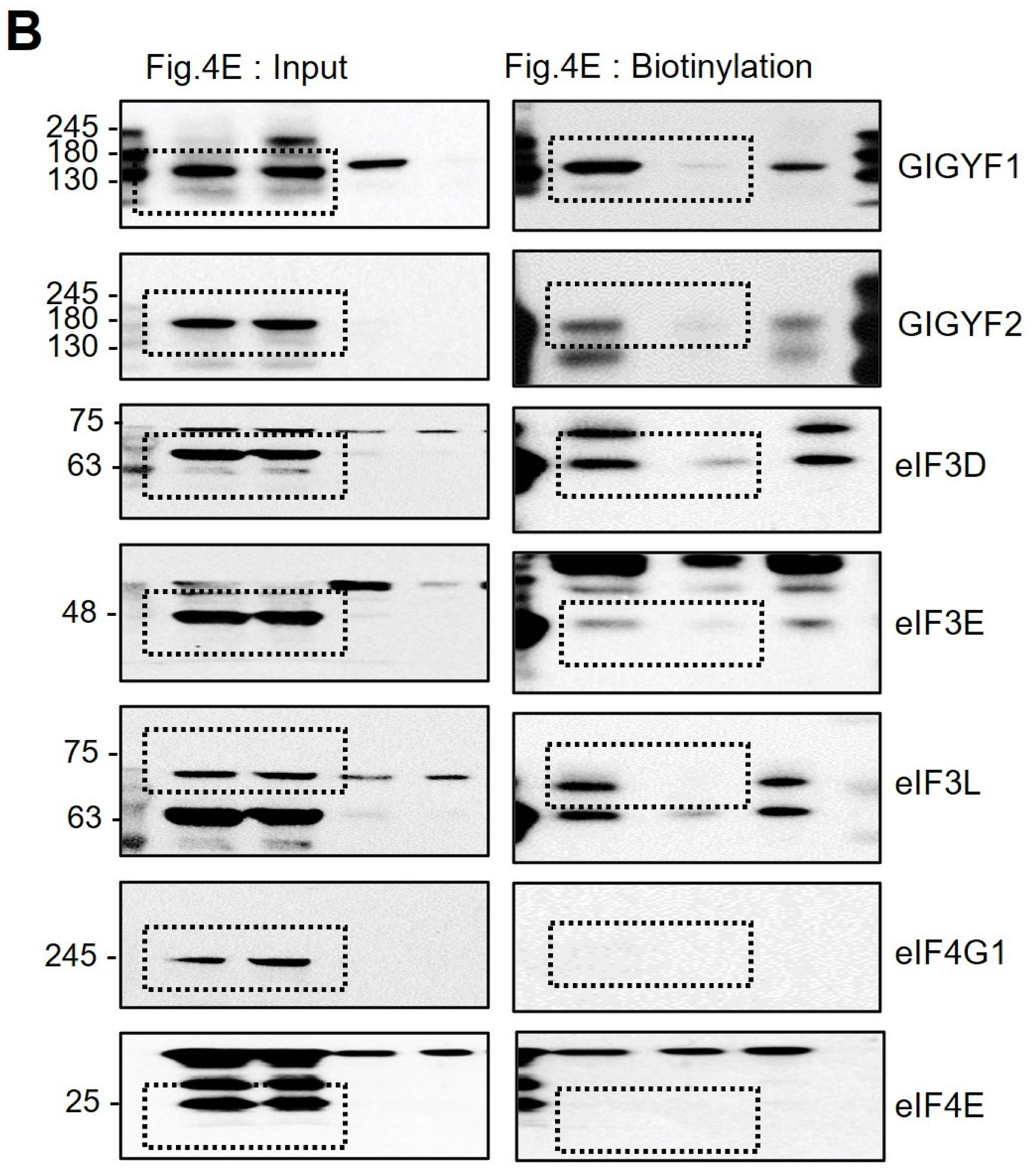


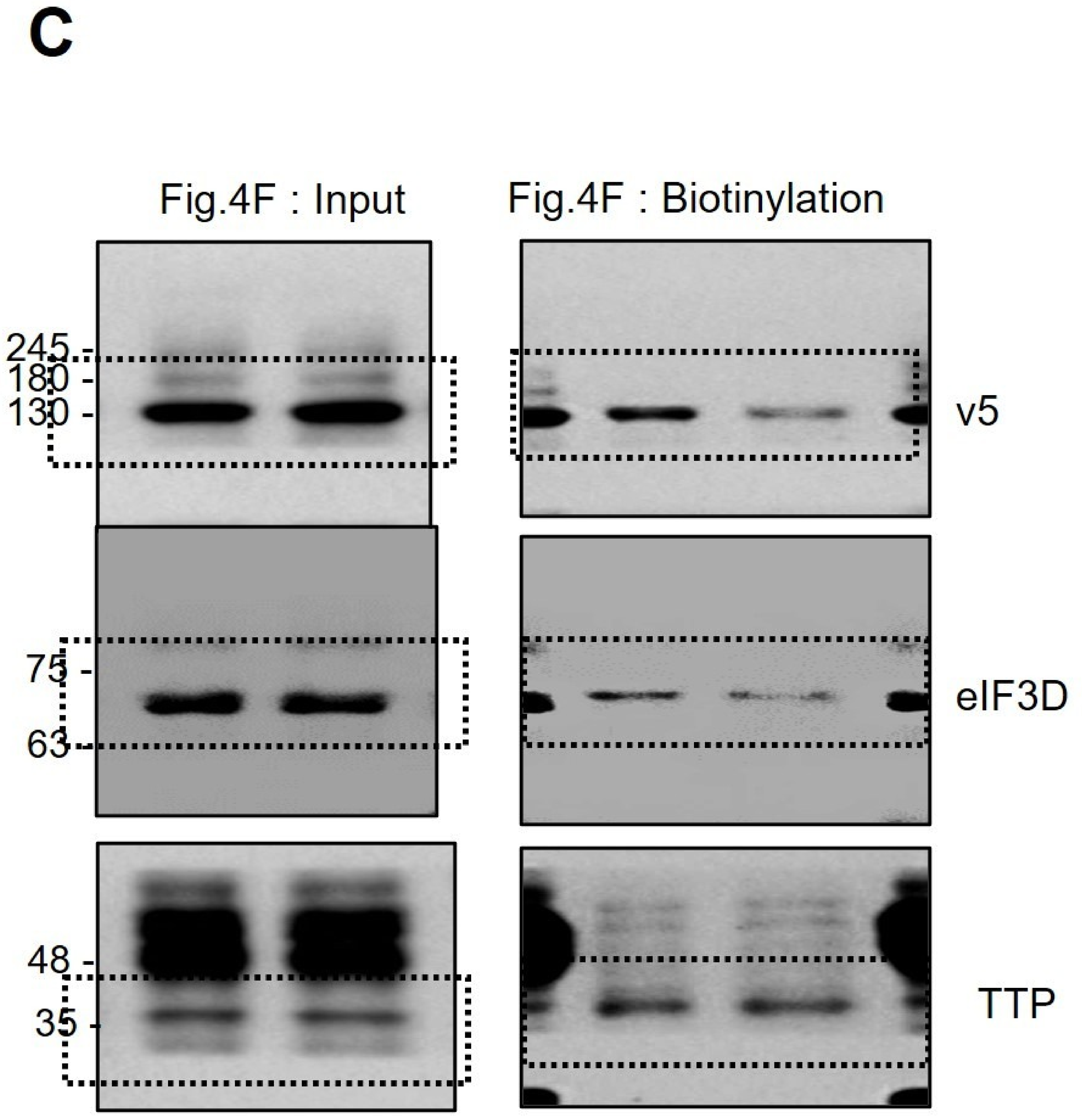


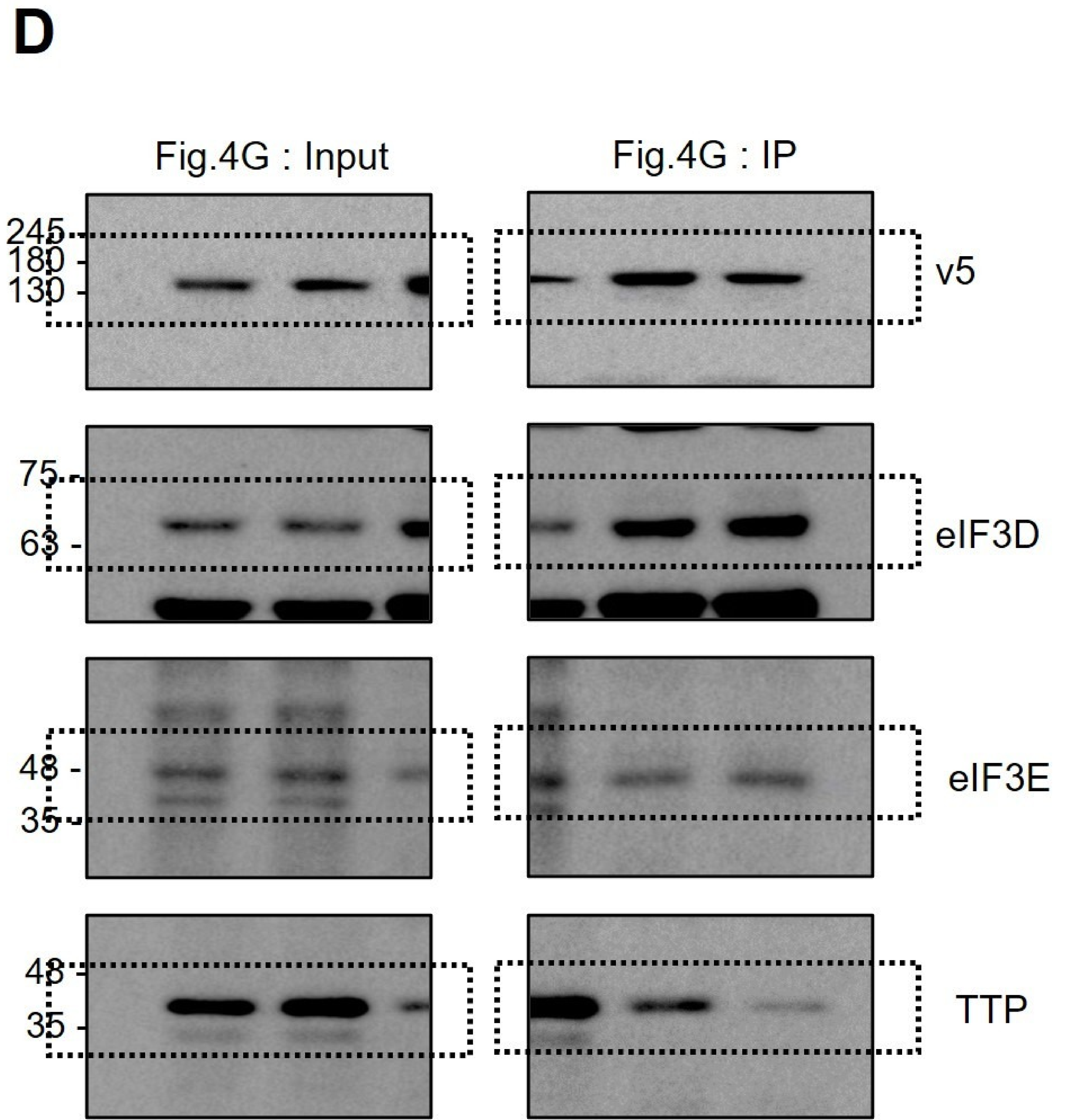

### Supplementary Figure. 7 ; Original blots for Supplementary Figure. 6

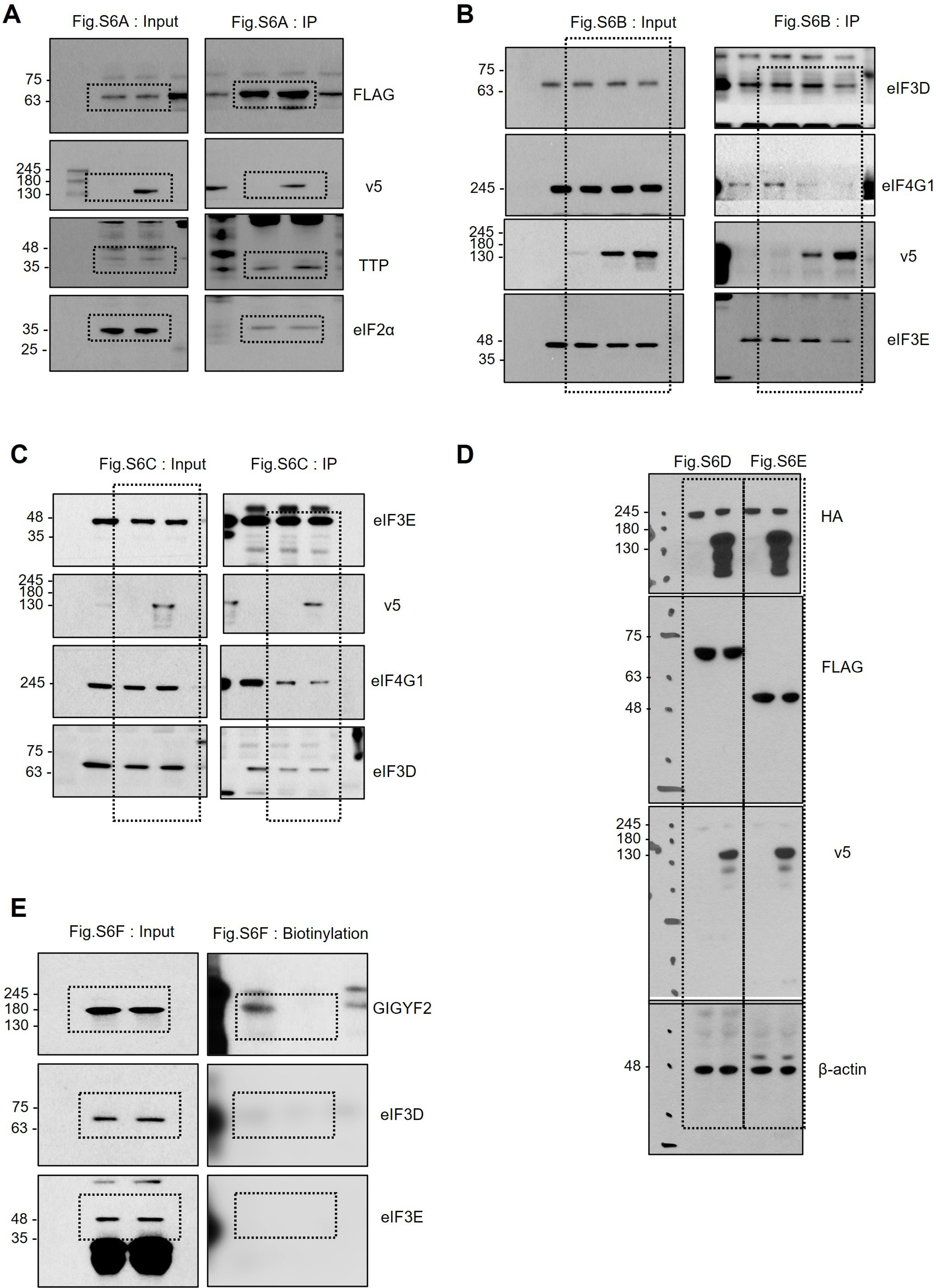

**F**

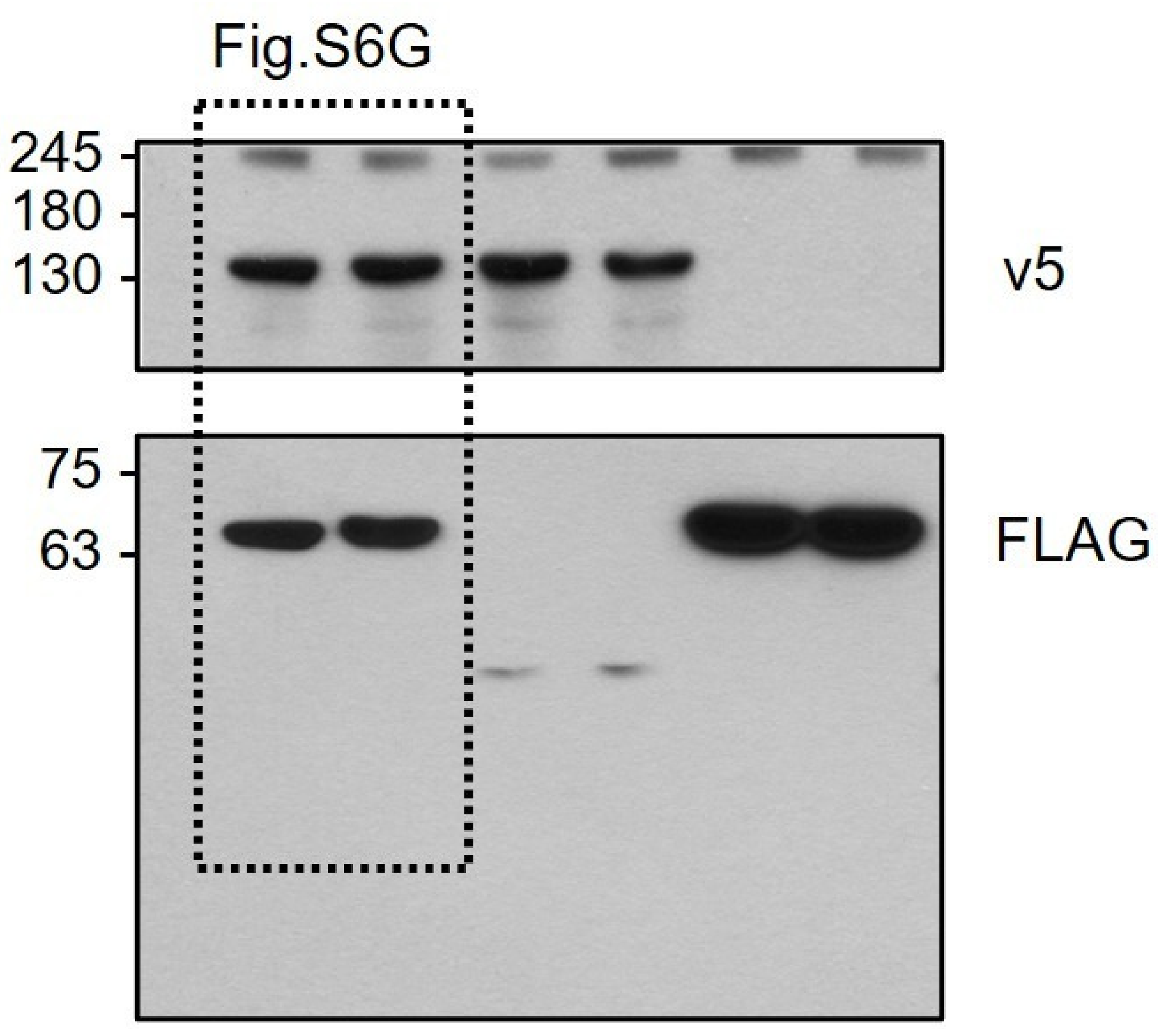
Fig.S6G
245
180
130
v5
75
63
FLAG

**G**

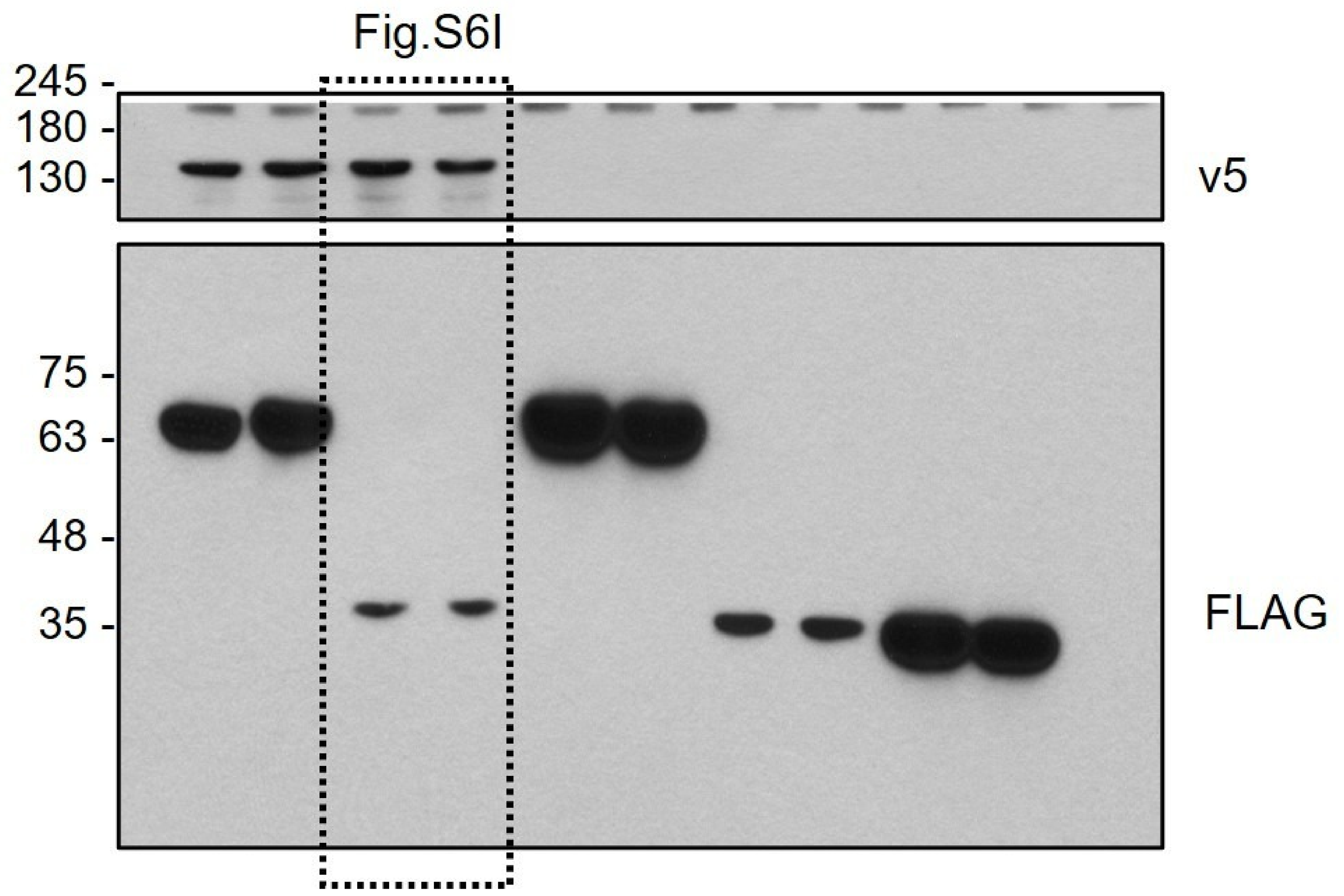
Fig.S6I
245
180
130
v5
75
63
48
35
FLAG

**H**

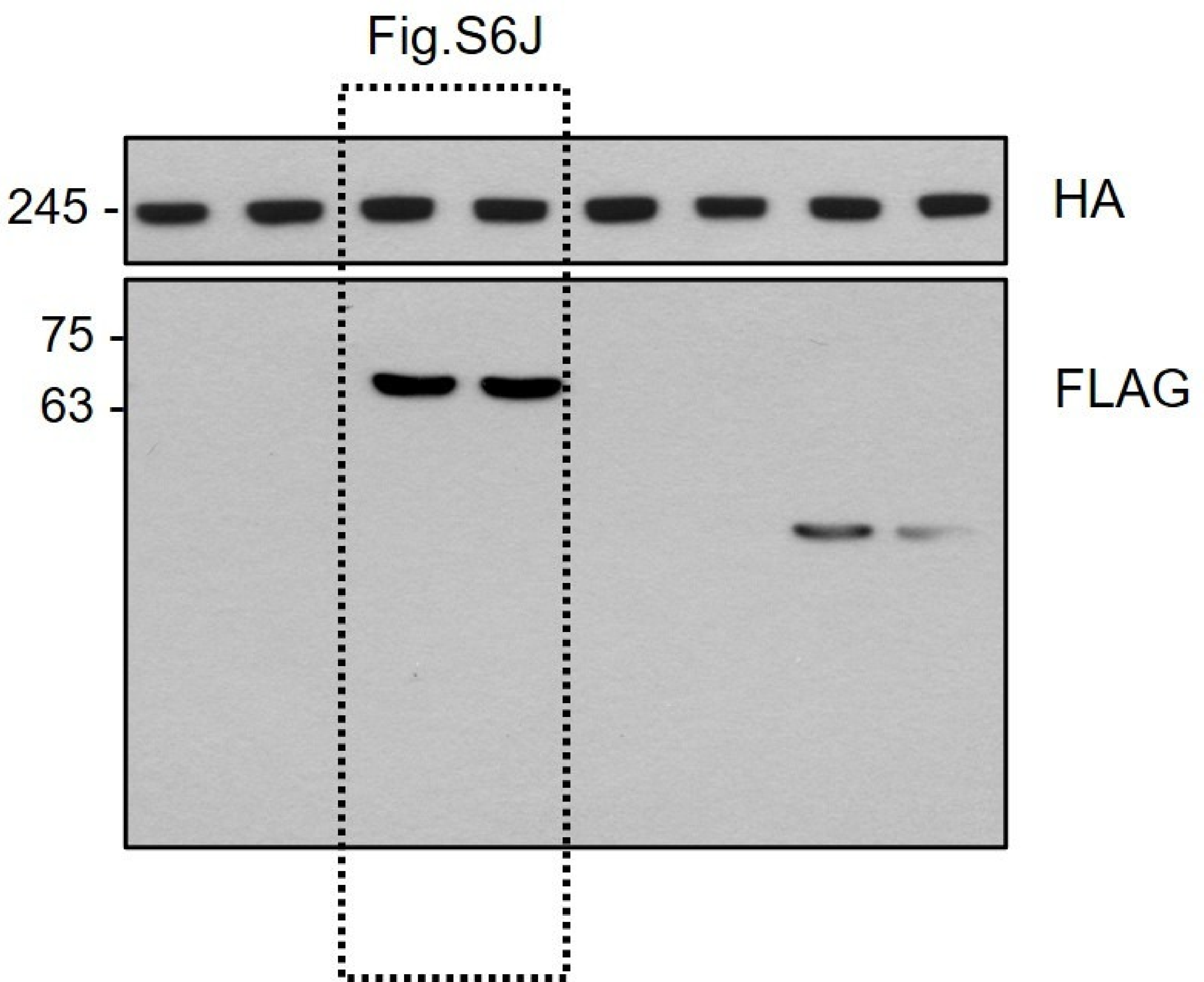
Fig.S6J
245
HA
75
63
FLAG
